# Unleashing the Potential of Noncanonical Amino Acid Biosynthesis for Creation of Cells with Site-Specific Tyrosine Sulfation

**DOI:** 10.1101/2022.03.25.485857

**Authors:** Yuda Chen, Shikai Jin, Mengxi Zhang, Kuan-lin Wu, Anna Chang, Shichao Wang, Zeru Tian, Peter G. Wolynes, Han Xiao

## Abstract

Incorporation of noncanonical amino acids (ncAAs) into proteins holds great promise for modulating the structure and function of those proteins and for influencing evolutionary dynamics in organisms. Despite significant progress in improving the efficiency of translational machinery needed for incorporating ncAAs, exogenous feeding of high concentrations of chemically-synthesized ncAAs, especially in the case of polar ncAAs, is required to ensure adequate intracellular ncAA levels. Here, we report the creation of autonomous cells, both prokaryotic and eukaryotic, with the ability to biosynthesize and genetically encode sulfotyrosine (sTyr), an important protein post-translational modification with low membrane permeability. We discovered the first enzyme catalyzing tyrosine sulfation, sulfotransferase 1C1 from *Nipponia nippon* (*Nn*SULT1C1), using a sequence similarity network (SSN). The unique specificity of *Nn*SULT1C1 for tyrosine has been systematically explored using both bioinformatics and computational methods. This *Nn*SULT1C1 was introduced into both bacterial and mammalian cells so as to yield organisms capable of biosynthesizing high levels of intracellular sTyr. These engineered cells produced site-specifically sulfated proteins at a higher yield than cells fed exogenously even with the highest level of sTyr reported in literature. We have used these autonomous cells to prepare highly potent thrombin inhibitors with site-specific sulfation. By enhancing ncAA incorporation efficiency, this added ability of cells to biosynthesize ncAAs and genetically incorporate them into proteins greatly extends the utility of genetic code expansion methods.

**TOC:** 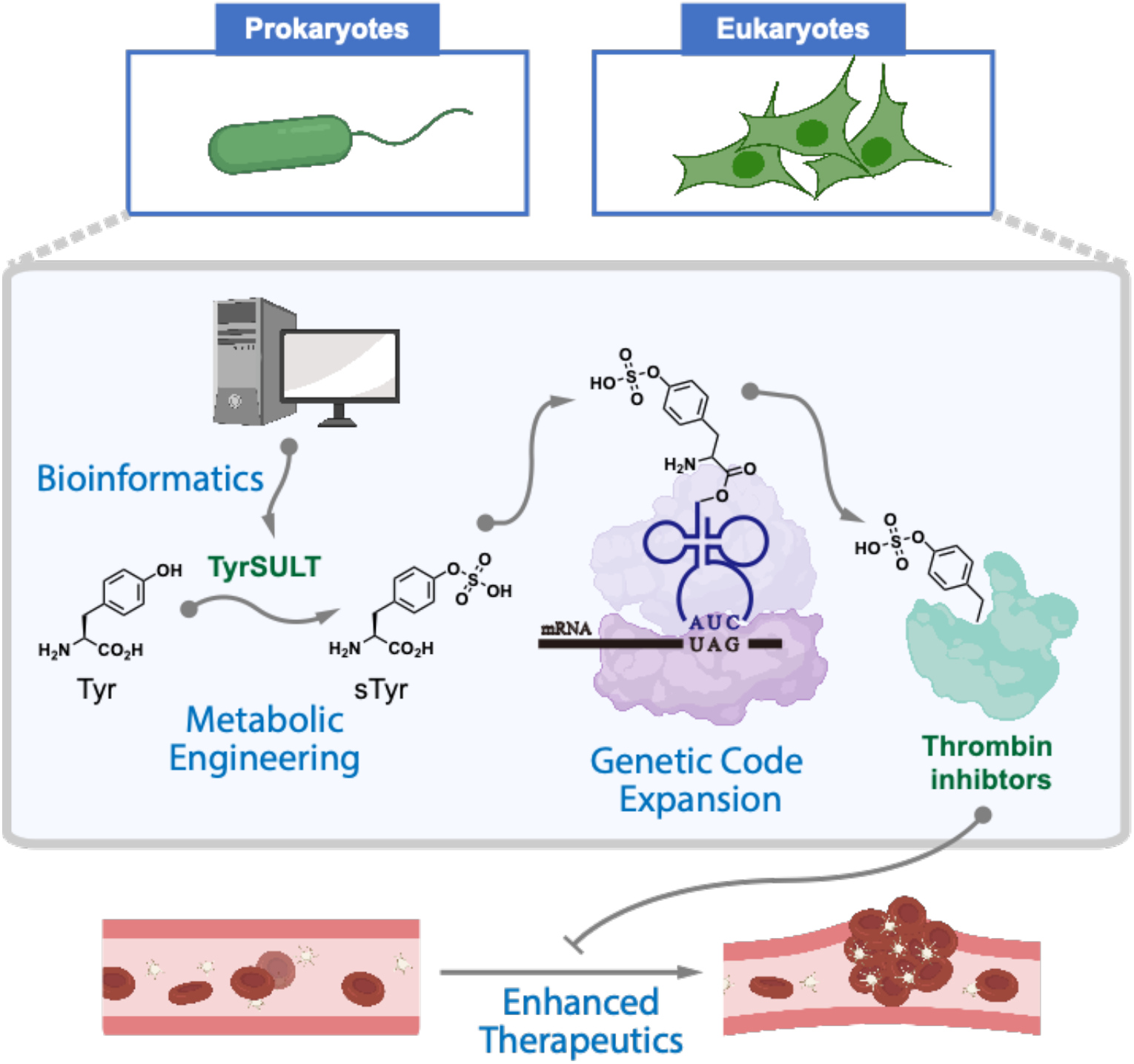

---

With the rare exceptions of pyrrolysine and selenocysteine, a standard set of 20 amino acid building blocks, containing a limited number of functional groups, is used by almost all organisms for the biosynthesis of proteins. The use of Genetic Code Expansion technology to enable the site-specific incorporation of noncanonical amino acids (ncAAs) into proteins in living cells has transformed our ability to study biological processes and develop modern medicines.^1–6^ The genetic encoding of ncAAs with distinct chemical, biological, and physical properties requires the engineering of bioorthogonal translational machinery, consisting of an evolved aminoacyl-tRNA synthetase/tRNA pair and a “blank” codon.^1,7,8^ The high intracellular concentration of ncAA required to render this machinery operative has usually been achieved via chemical synthesis of the ncAA and its exogenous addition at high levels to the cell culture medium. However, unlike canonical amino acids, ncAAs lack specific and efficient transporters that enable cellular internalization. Furthermore, lipid bilayers are impermeable to most polar and charged ncAAs.^9–13^ These factors greatly limit the efficiency of ncAA incorporation into proteins using the Genetic Code Expansion technology.^9,11–15^

Strategies for engineering the structures of ncAAs or ncAA-binding proteins have been employed to improve the cellular uptake of ncAAs. In 2017, the Schultz group adopted a dipeptide strategy to enable the cellular uptake of phosphotyrosine. The phosphotyrosine-containing dipeptide can be synthesized and transported into cells via an adenosine triphosphate (ATP)-binding cassette transporter, followed by hydrolysis of the dipeptide by nonspecific intracellular peptidases.^9^ In the same year, the Wang lab developed a two-step strategy for producing proteins with site-specific tyrosine phosphorylation.^10^ This strategy utilized the incorporation of a phosphotyrosine anologue with a cage group, followed by chemical deprotection of the purified proteins. But the synthesis and purification of these dipeptides are challenging, and the required post-purification treatments limit the applicability of this methodology to efficient incorporation of phosphotyrosine into proteins. As an alternative approach, periplasmic binding proteins (PBPs) have been engineered to have improved affinities for specific ncAAs.^15^ These mutant PBPs enhanced uptake of the respective ncAAs up to 5-fold, as evidenced by elevated intracellular ncAA concentrations and the yield of ncAA-containing green fluorescent proteins.^15^ Nevertheless, the engineered PBP species are only applicable to a subset of ncAAs, and exogenous feeding of high concentrations of the ncAAs is still required. The problem of ncAA uptake could potentially be bypassed by intracellular biosynthesis of the ncAAs.^14,16–19^ For example, phosphothreonine (pThr) cannot be detected intracellularly even when cells are incubated with 1 mM pThr.^14^ The Chin group overcame the membrane impermeability of pThr by introducing the *Salmonella enterica* kinase, PduX, which converts *L-*threonine to pThr intracellularly.^14^ This biosynthesis of pThr generated intracellular pThr at levels greater than 1 mM, sufficient for genetic incorporation of this amino acid.^14^ A similar strategy was recently applied to the creation of autonomous bacterial cells that can biosynthesize and genetically incorporate *p*-amino-phenylalanine (*p*AF), 5-hydroxyl-tryptophan (5HTP) and dihydroxyphenylalanine (DOPA), although no autonomous eukaryotic cells have been reported.^17,19,20^ We see there that additional biosynthetic pathways for producing polar or negatively-charged ncAAs would greatly expand the utility of genetic code expansion methods.

Tyrosine sulfation is an important post-translational modification of proteins that is important for a variety of biomolecular interactions, including chemotaxis, viral infection, anti-coagulation, cell adhesion, and plant immunity.^21–28^ Despite its importance and ubiquity, protein sulfation has been difficult to study due to the lack of general methods for preparing proteins with defined sulfated residues.^25,29^ To circumvent this challenge, efforts have been previously made to site-specifically incorporate sulfotyrosine (sTyr) using the Genetic Code Expansion technology.^30^ The resulting sTyr incorporation systems have enabled several applications, including generation of therapeutic proteins with defined sulfated tyrosines, evolution of sulfated anti-gp120 antibodies, and confirmation of tyrosine sulfation sites.^29,31–34^ To achieve reasonsble expression levels of sulfated proteins in *E. coli*, however, most studies have required the exogenous feeding of 3-20 mM sTyr to compensate for low intracellular uptake of extracellular sTyr.^31,34^

Here, we report the generation of completely autonomous prokaryotic and eukaryotic organisms capable of incorporating sTyr into proteins (**Fig. 1A**). sTyr is biosynthesized using a new sulfotransferase discovered using a sequence similarity network (SSN). sTyr is subsequently incorporated into proteins in response to a repurposed stop codon. The molecular properties of this new sulfotransferase were explored using bioinformatics and computational approaches, revealing a loop structure and several residues in binding pocket within this enzyme responsible for its unique specificity for tyrosine. The further optimization of the genome and sTyr biosynthetic pathway of both prokaryotic and eukaryotic cells leads to greater expression yields of sulfated proteins than experienced with cells exogenously fed with sTyr. The utility of these sTyr autonomous cells is demonstrated by using them to produce highly potent thrombin inhibitors.

**Figure 1:**
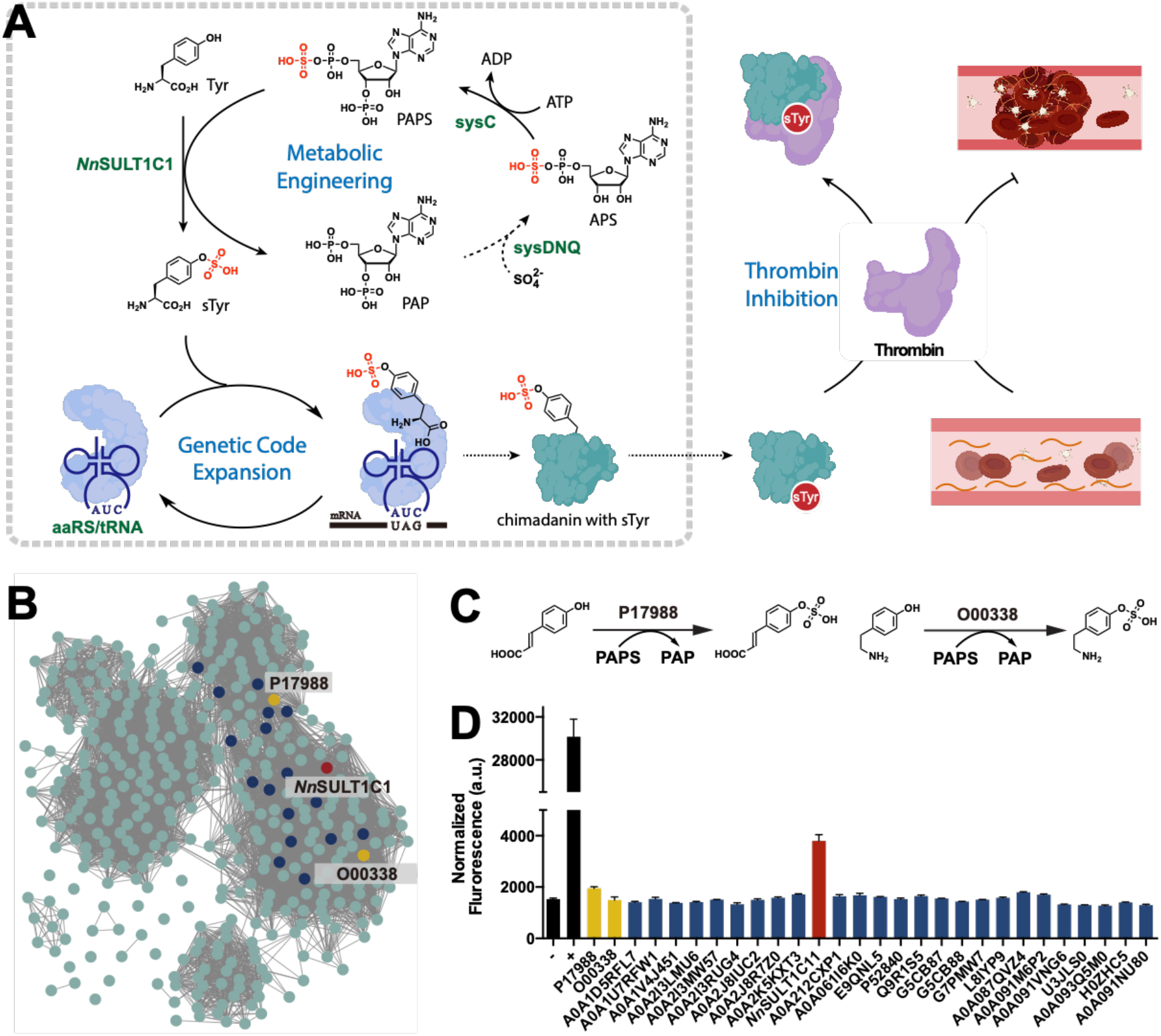
Discovery of tyrosine sulfotransferase from sequence similar network. **(A)** sTyr was biosynthesized from tyrosine and PAPS in the presence of sulfotransferase identified in this study. The resulting biosynthesized sTyr was site-specifically incorporated into thrombin inhibitors, yielding enhanced thrombin inhibition **(B)** Sequence similar network (SSN) generated by EFI-EST server with *Rn*SULT1A1 as an input sequence and E-value of 5. Each circle stands for a representative node containing sequences with over 80% identity. Edges detection threshold was set at an alignment score of 110. The upper and lower yellow representative nodes are *Rn*SULT1A1 (P17988) and *Hs*SULT1C2 (O00338), respectively. **(C)** Schematic representation of reported sulfation reactions of P17988 and O00338. **(D)** Screening of tyrosine sulfotransferases with green fluorescent protein assay. All tested proteins are included in the representative nodes of (B) and *Nn*SULT1C1 is the protein with red label (B).

## Results

### Discovery of Tyrosine Sulfotransferase using a Sequence Similarity Network

In nature, sulfotransferases allow many organisms to utilize an active form of sulfate, 3’-phosphoadenosine-5’-phosphosulfate (PAPS), for biosynthetic purposes.^35,36^ Based on their substrate preference and cellular location, sulfotransferases can be grouped into three major families, tyrosylprotein sulfotransferase (TPST), cytosolic sulfotransferase (SULT), and carbohydrate sulfotransferase (CHST).^37,38^ To identify the enzyme responsible for sulfation of cytoplasmic tyrosine, we focused on cytosolic sulfotransferases. These enzymes catalyze sulfation of a wide variety of endogenous compounds, including hormones, neurotransmitters, and xenobiotics.^37^ Based on their reported substrate specificities, we first examined SULT1A1 and SULT1A3 from *Homo sapiens*, SULT1A1 from *Rattus norvegicus*, and SULT1C1 from *Gallus gallus*,^37,39^ all of which are known to recognize multiple phenolic substrates. To explore the activity of these sulfotransferases towards tyrosine, we used a green fluorescent protein assay.^17,40^ These four sulfotransferase genes were codon-optimized for *Escherichia coli* and cloned into the pBad vector. To generate a suppression plasmid for sTyr incorporation, we used pUltra-sTyr plasmid encoding the engineered *Methanococcus jannaschii* tyrosyl-tRNA synthetase (sTyrRS) and its corresponding *Mj*tRNA^Tyr^_CUA_.^30,34^ The suppressor plasmid (pUltra-sTyr) was used to suppress the amber codon (Asp134TAG) within a sfGFP variant encoded by the pLei-sfGFP134TAG plasmid in the presence of sTyr. Expression of full-length sfGFP was carried out in LB medium for 16 hours in parallel with controls BL21(DE3) harboring pUltra-sTyr, pLei-sfGFP134TAG and pBad-Empty in the presence and absence of exogenously fed 1 mM sTyr. As expected, sfGFP was expressed in the presence of 1 mM sTyr fed in controls cells (**Fig. S1**). Unfortunately, none of these four sulfotransferases led to sfGFP expression, indicating the failure of the biosynthesis of sTyr. To circumvent the limited substrate range of the reported sulfotransferases, we accessed the full repertoire of protein sequence diversity in nature by using a sequence similarity network (SSN, **Fig. 1B**).^41^ SSNs provide an effective way to visualize and analyze the relatedness of massive protein sequences on the basis of similarity thresholds of amino acid sequence.^42^ We initially created an SSN with EFI-ESI based on rat SULT1A1 as an input sequence, since its cognate substrate *p*-coumaric acid is similar to tyrosine (**Fig. 1B** and **1C**).^39^ An alignment score of 110 was set to limit the edges and a sequence identity of 80% was used to generate representative nodes, which resulted in a final SSN of 391 representative enzyme sequences. Interestingly, we found that human SULT1C2, whose substrate is tyramine, was in a different cluster of the SSN (**Fig. 1B** and **1C**).^37^ We hypothesized that enzymes with high sequence similarity to rat SULT1A1 and human SULT1C2 would possess the ability to carry out the sulfation of tyrosine. To test this hypothesis, we selected 27 sequences from the SSN based on their proximity to both *Rn*SULT1A1 and *Hs*SULT1C2. These selected genes were cloned into the pBad vector and tested with the green fluorescent protein assay. To our delight, a 2.5-fold increase in fluorescence was observed for cells expressing A0A091VQH7 compared to cells not given exogenous sTyr, suggesting that sTyr was biosynthesized intracellularly and incorporated into sfGFP proteins (**Fig. 1D**). A0A091VQH7 is a putative sulfotransferase from *Nipponia nippon*, with over 90% sequence indentity with SULT1C1 reported in other species. Thus, we name A0A091VQH7 as *Nn*SULT1C1 hereafter.^43^

### Molecular Basis of *Nn*SULT1C1 Action in the Sulfation of Tyrosine

To explore the origin of the unique tyrosine specificity of *Nn*SULT1C1 among all the sulfotransferases tested, we first analyzed the phylogenetic relationships of the enzymes. Sulfotransferase amino acid sequences were used to generate a phylogenetic tree using the unweighted pair group method with arithmetic mean (UPGMA) by MEGA X software package (**Fig. S2**).^44^ The tree is subdivided into three major subfamilies, among which *Nn*SULT1C1 falls into subfamily I containing bird sulfotransferases. Most sequences from subfamilies II and III are derived from rodent and primate groups, respectively. To further analyze the molecular basis of the unique tyrosine specificity of *Nn*SULT1C1, we performed a multiple sequence alignment (MSA) of all sequences within subfamily I of the phylogenetic tree (**Fig. S3**). This sequence alignment revealed that most regions of *Nn*SULT1C1, including the PAPS-binding site, are highly conserved throughout the tree except for a highly variable region corresponding to *Nn*SULT1C1 residues 94-103 (SIQEPPAASY) and residues likely involved in substrate binding pocket.^45,46^ To explore the contribution of this highly variable region and these residues to substrate binding, the structure of *Nn*SULT1C1 was predicted via Alphafold 2. Alphafold 2 is a machine learning approach that has been sbown to predict protein structure with a high degree of accuracy.^47–50^ More than 90% of the residues in the predicted *Nn*SULT1C1 structure show Local Distance Difference Test (IDDT) values over 90, indicating they have a significant likelihood of very high accuracy in the predicted structure. Similar to the structures of other cytosolic sulfotransferases, the overall predicted structure of *Nn*SULT1C1 is composed of classical α/β motifs (**Fig. 2A**).^51,52^ This structure includes a β sheet surrounded by α-helices, giving rise to a narrow substrate-binding site (**Fig. 2A**).^53^ We found that the highly variable region (94-102 residue) of *Nn*SULT1C1 constitutes a loop for the substrate entry, which also aligns with the substrate entry loop of human cytosolic sulfotransferase **(Fig. S4)**.^37^ The deletion of this loop on *Nn*SULT1C1, however, only results in 22% decrease of its activity to produce fluorescent protein with sTyr. (**Fig. 2B)**

**Figure 2:**
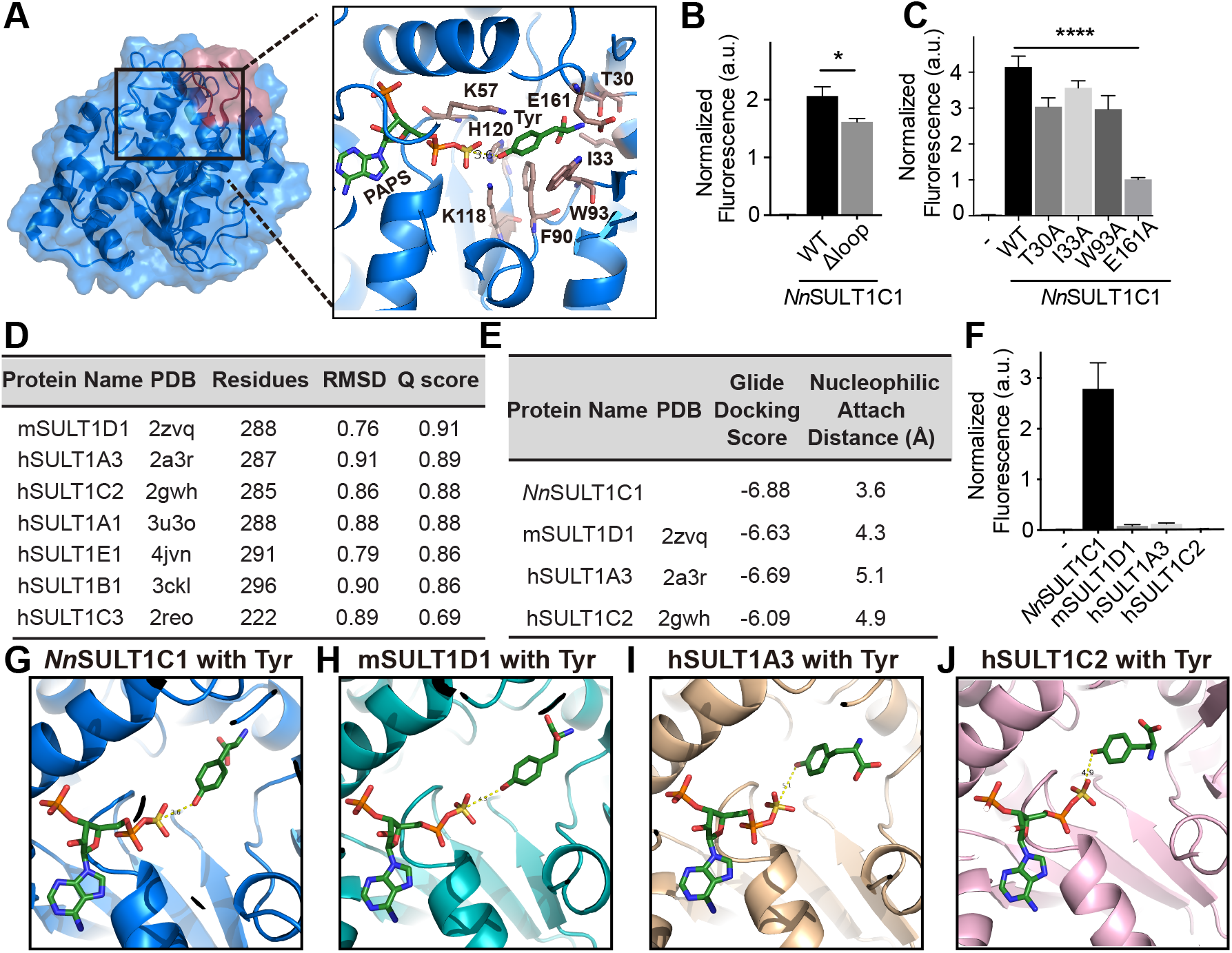
Exploring the mechanism of unique tyrosine specificity of NnSULT1C1. **(A)** *Nn*SULT1C1 structure (blue) predicted by AlphaFold2 and its active site consisting PAPS and Tyr. Tyr was docked into *Nn*SULT1C1 containing PAPS by Glide v8.1 in Schrödinger software. **(B)** Green fluorescent protein assay with wildtype *Nn*SULT1C1 (wt) or *Nn*SULT1C1 without the SIQEPPAASY (Δloop). **(C)** Green fluorescent protein assay with wildtype *Nn*SULT1C1 (wt) or *Nn*SULT1C1 with alanine mutation at indicated resiudes. **(D)** Structural similarity search of *Nn*SULT1C1 using the PDBeFold web server. **(E)** Characterization of Tyr docking with *Nn*SULT1C1 and its structurally similar sulfotransferases via docking score and nucleophilic attack distance. Docking scores were calculated using Glide v8.1 in Schrödinger software. Nucleophilic attack distance was defined as the distances between Tyr phenolic alcohol and PAPS sulfonate. **(F)** Comparison of tyrosine sulfation activity of *Nn*SULT1C1 and its structurally similar sulfotransferases using green fluorescent protein assay. Cells without any sulfotransferase (-) were used as control. **(G-J)** Tyr docking position with *Nn*SULT1C1 **(G)**, mSULT1D1 **(H)**, hSULT1A3 **(I)**, and hSULT1C2 **(J)**. PAPS and Tyr are shown as sticks with green carbon. Docking was performed by Glide v8.1 in Schrödinger software with the same parameters in (A). \**p*<0.05; \*\*\*\**p*<0.0001.

To further explore the other residues involved in substrate binding of *Nn*SULT1C1, we performed protein-ligand docking using Glide v8.1 in Schrödinger software package v2018.4.^54^ The Tyr was docked to the *Nn*SULT1C1 using OPLS_3 force field and the lowest energy pose was monitored.^55^ For each docking experiment, 200 maximum output poses for each protein were set and Emodel energy was used for ranking the top 50 poses. The docking structure suggests that the α-amino group of Tyr is stabilized by *Nn*SULT1C1 residues Glu161, Thr30, Ile33, and Trp93. The π-π stacking interactions between Tyr and Phe90 are likely to improve the packing interaction (**Fig. 2A**). The phenolic hydroxy group of Tyr is in the proper Lys-Lys-His catalytic site to engage in sulfuryl transfer. The His120 residue serves as a catalytic base that can remove the proton from Tyr and stabilize the transition state. The Lys57 and Lys118 residues interact with and stabilize the sulfuryl group of PAPS and the phenolic hydroxy group of Tyr, respectively. To validate the the contribution of these residues interacting with Tyr on *Nn*SULT1C1 activity, Thr30, Ile33, Trp93, and Glu161 were mutated to alanine separately. Alanine mutation at Thr30, Trp93, or Glu161 significantly decreased the activity of *Nn*SULT1C1 **(Fig. 2C)**. Among these residues, the E161A mutation exhibits the largest decrease in activity, confirming its important interaction with Tyr. To further explore whether other sulfatransferases may also carry out the tyrosine sulfation, we performed a structure similarity search using the PDBeFold. Based on the Q score, the three proteins with structures most similar to *Nn*SULT1C1 are mouse SULT1D1 (pdb: 2zvq), human SULT1A3 (pdb: 2a3r) and human SULT1C2 (pdb: 2gwh, **Fig. 2D**). The overall secondary structure of *Nn*SULT1C1 alignmented well with 2zvq, which indicates its structural consistency with the other cytosolic sulfotransferases. (**Fig. S5)** To further illustrate the unique specificity of *Nn*SULT1C1 for Tyr, dockings of Tyr to the most similar sulfotransferases, including mSULT1D1, hSULT1A3, and hSULT1C2, were carried out using Glide v8.1 in the Schrödinger software package v2018.4 following the same method of Tyr docking used for *Nn*SULT1C1.^54^ Docking of Tyr to *Nn*SULT1C1 exhibits the lowest Glide Docking score of -6.88 and the closest distance between the phenolic hydroxyl group and PAPS sulfonate. **(Fig. 2E**) This result is consistent with the optimal ability of *Nn*SULT1C1, to generate sTyr-containing sfGFP in the green fluorescent protein assay among all tested sulfotransferases **(Fig. 2F**). The key step of the sulfotransfer reaction involves an S_N_2-type nucleophilic attack on the PAPS sulfonate by the phenoxide of Tyr. Compared with mSULT1D1, hSULT1A3, and hSULT1C1, the docking of Tyr in *Nn*SULT1C1 results in the closest distance (3.6Å) between the sulfur atom of PAPS and the phenolic hydroxyl group **(Fig. 2G-J**). Furthermore, the acceptor phenolic hydroxyl group of Tyr lies on the backside of the S-O bond of PAPS in the Tyr docking structure with *Nn*SULT1C1, indicating a more proper orientation for the nucleophilic attack **(Fig. 2G**).

### Biosynthesis and Genetic Encoding of sTyr in *Escherichia coli*

Having identified *Nn*SULT1C1 as a functional tyrosine sulfotransferase, we explored whether the biosynthesized sTyr can be genetically incorporated into proteins in *E. coli* in response to the amber codon. As an initial goal, we wanted to increase sTyr production in these cells in order to optimize its availability for incorporation into proteins. Since *Nn*SULT1C1 utilizes tyrosine and PAPS for producing sTyr, we quantified sTyr production in five knockout *E. coli* cell lines in which the gene knockout has been shown to improve the yield of either tyrosine or PAPS in *E. coli*.^56–59^ To evaluate the effect of knocking out these genes on the biosynthesis of sTyr, we transformed the suppression plasmid pUltra-sTyr, reporter plasmid pET22b-T5-sfGFP151TAG, and the biosynthesis plasmid pEvol-*Nn*SULT1C1 into wild type *E. coli* BW25113 or knockout strains (**Fig. 3A**). The expression of sfGFP with sTyr at position 151 (sfGFP-sTyr) was carried out in LB medium for 18 h. To our delight, we found that knockout of the cysH gene significantly improved the production of sTyr-containing sfGFP, compared to that seen in the wildtype BW25113 strain (**Fig. 3B**). CysH encodes the PAPS sulfotransferase responsible for degradation of PAPS to 3’-phosphoadenosine-5’-phosphate (PAP). This observation of enhanced sfGFP-sTyr production in BW25113ΔcysH is consistent with the previous report that knockout of the cysH gene can increase cellular PAPS concentration and the production of sulfated products in *E. coli*.^59,60^ Next, we examined whether manipulation of PAPS synthetic and recycling pathways in *E. coli* could enhance intracellular PAPS levels. We first amplified the gene cysDNC encoding adenosine-5’-triphosphate (ATP) sulfurylase and adenosine 5’-phosphosulfate (APS) kinase to increase the intracellular level of PAPS, followed by the introduction of the gene cycQ encoding adenosine-3′,5′-diphosphate (PAP) nucleotidase for PAP recycling.^61,62,39,60^ We found that cells with both PAPS synthesis and recycling pathways exhibited the largest increase in fluorescence, suggesting a higher expression level of sfGFP containing sTyr (**Fig. 3C**). To examine the contribution of the biosynthetic pathway to intracellular sTyr concentration, we measured the intracellular sTyr concentrations in cells when sTyr was either biosynthesized or delivered via exogenous feeding. Since a high concentration of exogenous ncAA has been shown to be associated with higher protein production using the Genetic Code Expansion technology, we screened the effect of exogenous sTyr concentrations to up to 27 mM, higher than all reported concentration added externally.^29–34^ Surprisingly, the cellular concentration of sTyr in cells endowed with the sTyr biosynthetic pathway is 756.3 μM, which is 28 fold higher than that from cells exogenously fed with 1 mM sTyr and higher even than in cells fed with 27 mM (**Fig. 3D**). Consistent with these intracellular levels of sTyr, in the context of Genetic Code Expansion technology, endogenous biosynthesis of sTyr results in much higher sfGFP-sTyr expression than that produced via exogenous feeding (**Fig. 3E**). Predictably, the *Nn*SULT1C1 expression level also has a significant influence on the production of sfGFP-sTyr. Since we found that *Nn*SULT1C1 expression induced by 15 mg/L *L*-arabinose yielded the highest production of sfGFP-sTyr, 15 mg/L *L*-ara was consistently used for induction of the sTyr biosynthetic pathway (**Fig. 3E**). We also screened other variables for sfGFP-sTyr expression, including expression medium, Tyr addition, SO_4_^2-^ addition, glycerol addition, which did not optimize the expression of sfGFP-sTyr. (**Fig. S6**) To further investigate the efficiency and specificity of incorporation of biosynthesized sTyr in these autonomous *E. coli* cells, sfGFP-sTyr proteins derived from exogenously fed sY and from biosynthesized sY were purified by Ni^2+^-NTA affinity chromatography and characterized by SDS-PAGE and ESI-MS. Intact sfGFP was only expressed after exogenous sTyr feeding or after induction of sTyr biosynthesis. The yield of sfGFP-sTyr derived from biosynthetic sTyr is 5.67 mg/L sfGFP-sTyr under the optimal condition, compared with 1.5 mg/L sfGFP-sTyr produced by feeding with 1 mM exogenous sTyr (**Fig. 3F**). The mass of sfGFP-sTyr produced from biosynthetic sTyr was 27, 674 Da, which is in good agreement with the calculated mass. (**Fig. 3G** and **3H**).

**Figure 3:**
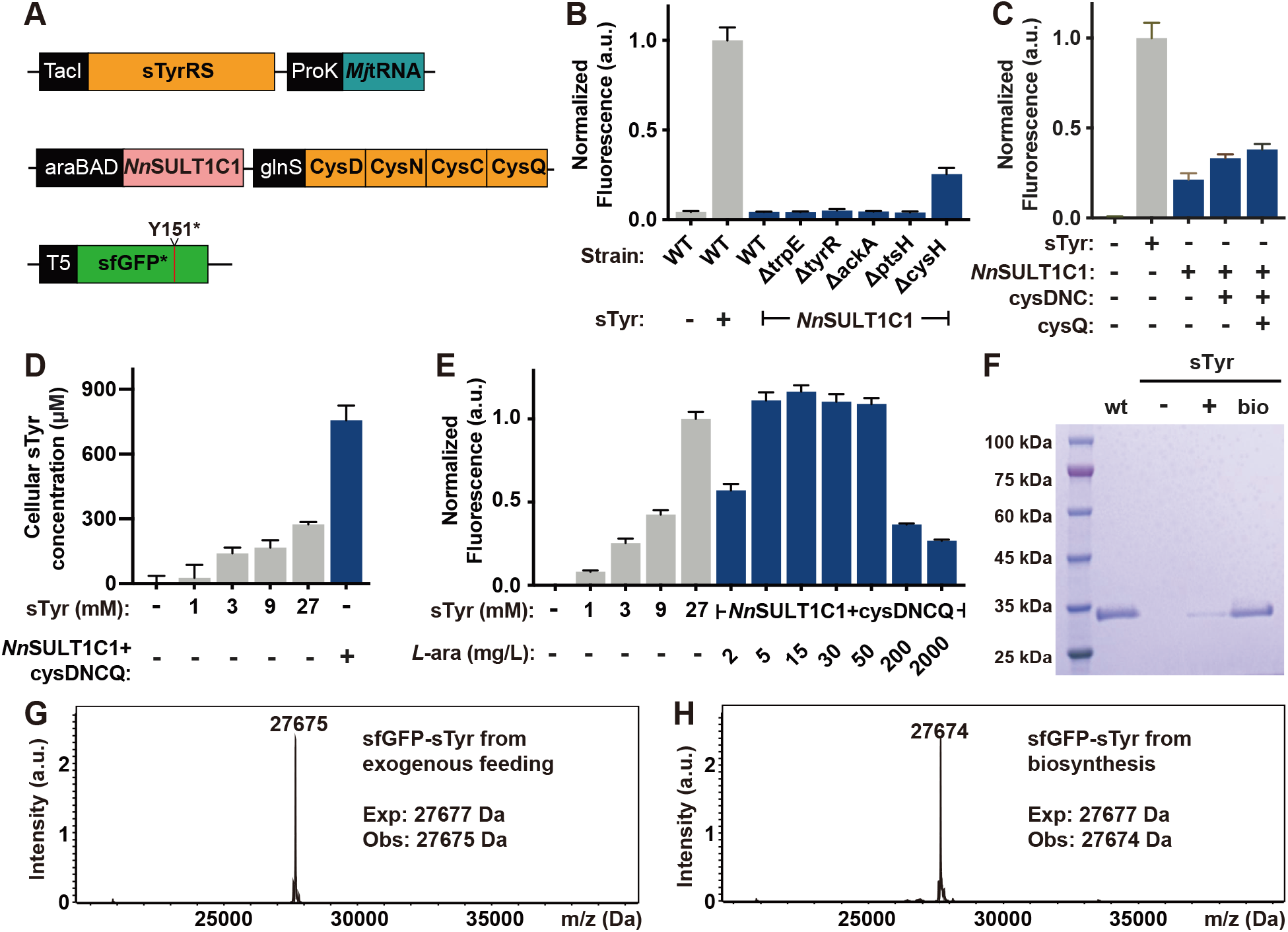
Generation of completely autonomous sTyr synthesizing *E. coli*. **(A)** Schematic representation of genetic circuits used for generating completely autonomous sTyr synthesizing *E. coli*. **(B)** Screening of the knockout strains for sfGFP-sTyr production after the expression of *Nn*SULT1C1. **(C)** The roles of PAPS recycling enzymes in producing sfGFP-sTyr using ΔcysH BW25113 strain. **(D)** Cellular concentrations of sTyr of cells with the addition of chemically synthesized sTyr or the biosynthesis of sTyr. **(E)** Production of sfGFP-sTyr from cells with the addition of chemically synthesized sTyr or the biosynthesized Tyr. The effect of *Nn*SULT1C1 expression level on producing sfGFP-sTyr was screened by altering the concentration of inducer, *l*-arabinose (*l*-ara). **(F)** SDS-PAGE analysis of sfGFPs expressed in LB in the presence (+) or absence (-) of exogenous 1 mM sTyr addition or when inducing *Nn*SULT1C1 expression (bio). (**G-H**) Mass spectra analysis of sfGFP-sTyr proteins expressed in cells with the addition of 1 mM chem ically synthesized sTyr or the biosynthesis of sTyr.

### Biosynthesis and Genetic Encoding of sTyr in mammalian cells

Post-translational tyrosine sulfation occurs exclusively in eukaryotes. Although this modification has been estimated to occur on 1% of all tyrosine residues in eukaryotic proteomes, its functional significance is not well understood.^35,63,64^ One approach to determining the biological importance of protein tyrosine sulfation is to express in living cells proteins that are sulfated in site-specific and homogeneous fashion, a goal that is difficult to achieve by chemical synthesis or recombinant expression. Genetic code expansion based on *E. coli*-derived tyrosyl-tRNA synthetase (*Ec*TyrRS)/tRNA has been proven to overcome these challenges by site-specifically incorporating sTyr in proteins in mammalian cells.^65,66^ To promote the efficient expression of mammalian proteins sulfated on specific tyrosines, we have created mammalian cells equipped with both sTyr biosynthetic and translational machinery. To generate mammalian cells capable of biosynthesizing sTyr, we used piggybac system to stably integrate *Nn*SULT1C1 into the genome of HEK293T cells, yielding the HEK293T-*Nn*SULT1C1 cell line (**Fig. 4A**).^67^ The *Ec*TyrRS/tRNA pair was used to construct pAcBac2.tR4-sTyrRS/EGFP*, containing EGFP with a stop codon at position 39 as well as two copies of *E. coli* and *Bacillus stearothermophilus* tRNA_CUA_^Tyr^ (**Fig. 4A**).^68^ To evaluate the function of *Nn*SULT1C1 in mammalian cells, pAcBac2.tR4-sTyrRS/EGFP* was transfected into HEK293T and HEK293T-*Nn*SULT1C1 cells, which were then incubated in the presence or absence of exogenous sTyr. The expression of EGFP was monitored by confocal microscopy at 2 days after transfection. As expected, the addition of 1 mM sTyr to HEK293T cells resulted in moderate expression of full-length EGFP, while minimal EGFP fluorescence was observed in the absence of sTyr addition (**Fig. 4B**). Gratifyingly, higher expression of EGFP was observed in HEK293T-*Nn*SULT1C1 cells without exogenous sTyr addition than that seen in HEK293T cells fed with 3 mM sTyr. In addition to confocal imaging, flow cytometry was used to quantify expression levels of EGFP in cells fed with exogenous sTyr and in cells biosynthesizing sTyr. As shown in Fig. 4C, significantly higher EGFP fluorescence was observed in HEK293T-*Nn*SULT1C1 cells endowed with sTyr biosynthetic capability than in HEK293T cells fed with 3 mM sTyr. As a direct evidence of sTyr biosynthesis in mammalian cells, cellular sTyr concentration in HEK293T-NnSULT1C1 is more than that in HEK293T cells fed with 3 mM sTyr. (**Fig. S7**) The fidelity of site-specific incorporation of sTyr was evaluated by mass spectral analysis of purified sTyr-containing EGFP proteins. The observed mass was 29761 Da, consistent with the calculated mass of EGFP with sTyr at position 39 and observed mass of EGFP39sTyr purified from HEK293T with external sTyr addition. (**Fig. 4D and S8**). These results demonstrate that the creation of mammalian cells autonomously able to biosynthesize sTyr and incorporate it into proteins significantly enhances the expression level of sTyr-containing protein in mammalian cells.

**Figure 4:**
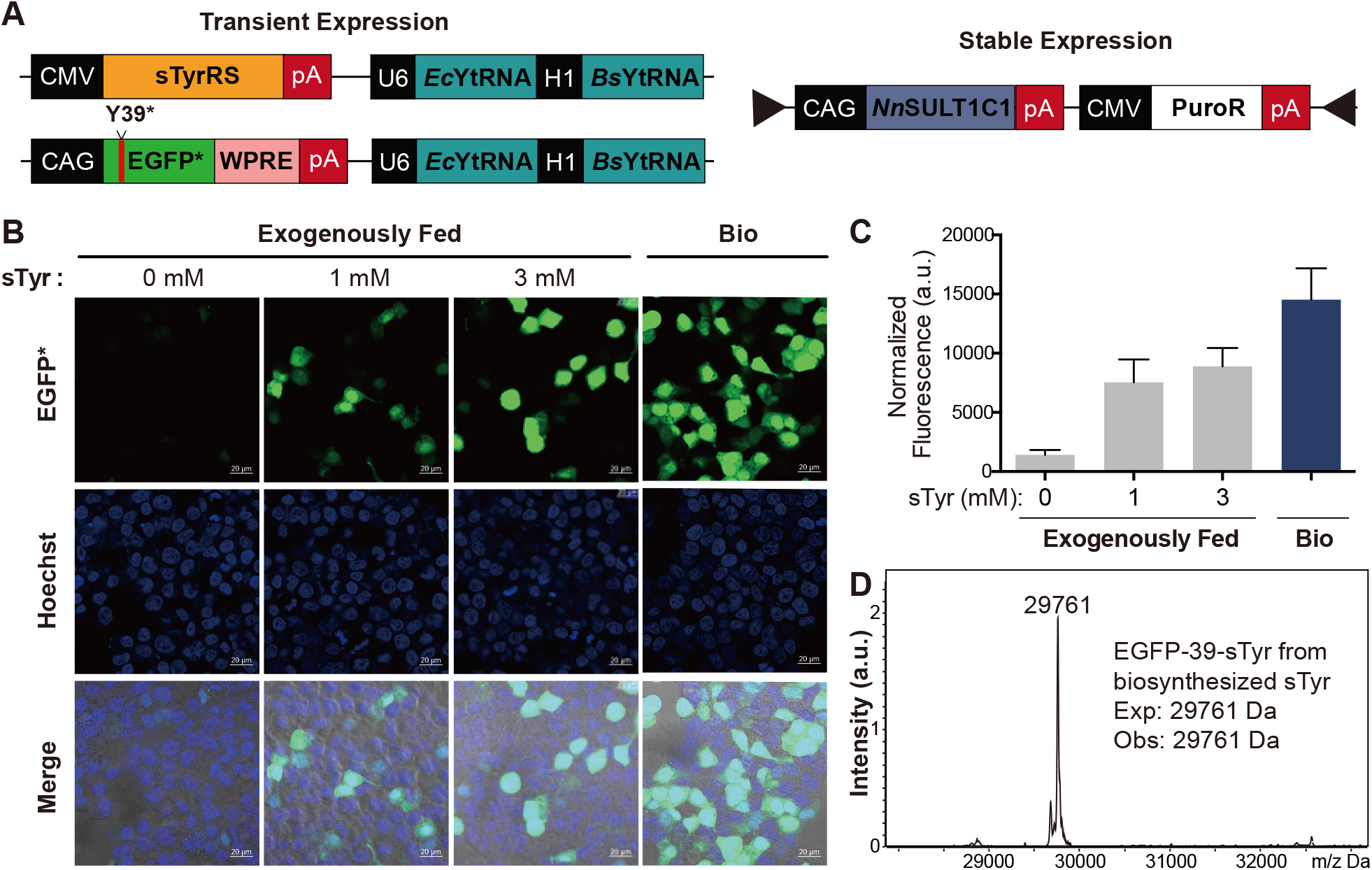
Generation of completely autonomous mammalian cells with sTyr-containing proteins. **(A)** Schematic representation of genetic circuits used for generating completely autonomous mammalian cells with sTyr-containing proteins. **(B)** Confocal images of HEK293T (exogenously fed) and HEK293T-*Nn*SULT1C1 (bio) cells expressing sTyrRS, tRNA_CUA_ and EGFP containing an amber codon at Tyr39 position. Scale bar = 20 μm. **(C)** Flow cytometric analysis of EGFP expression levels of HEK293T (exogenously fed) and HEK293T-*Nn*SULT1C1 (bio) cells with sTyrRS, tRNA_CUA_ and EGFP containing an amber codon at Tyr39 position. The normalized fluorescence was calculated by multiplying the geometric mean fluorescence by the percentage of EGFP-positive cells. Error bars represent standard deviations. **(D)** Mass spectra analysis of EGFP with sTyr (EGFP-39-sTyr) purified from HEK293T-*Nn*SULT1C1 cells.

### Using Completely Autonomous sTyr Biosynthetic Cells to Synthesize Potent Thrombin Inhibitors with Site-Specific Sulfation

Thrombin inhibitors represent an important class of anticoagulants used to prevent blood clotting. In addition, several thrombin inhibitors from hematophagous organisms have been shown to facilitate the acquisition and digestion of bloodmeal.^69–71^ Recent studies have reported that post-translational sulfation of these proteins has a dramatic effect on their inhibitory activity.^25,72^ For example, tyrosine sulfation of hirudin increases its affinity for thrombin more than 10 fold.^72,73^ Tyrosine sulfation of madanin-1 and chimadanin significantly increases their affinities for thrombin by promoting strong electrostatic interactions with positively-charged residues (**Fig. 5A**). Current methods for studying these site-specifically sulfated thrombin inhibitors rely heavily on solid-phase peptide synthesis and subsequent chemical ligation, processes that are time-consuming and may result in sub-optimal protein folding.^25,74,75^ To explore the generation sTyr-containing thrombin inhibitors using cells endowed with autonomous sTyr biosynthetic machinery, we chose both madanin-1 and chimadanin identified in the salivary gland of *haemaphysalis longicornis* (**Fig. 5B**).^25,76^ As shown in Figure 5A, sulfation of madanin-1 converts Tyr32 and Tyr35 to negative residues, thus enhancing madanin-1’s direct electrostatic interaction with the ε-amino groups of K236 and K240 located within the exosite II site of thrombin. To express the site-specifically sulfated thrombin inhibitors, we constructed plasmids encoding the thrombin inhibitor and substituted with amber codons at either or both of the indicated Tyr sites. sTyr-containing inhibitors were expressed by transforming ΔcysH BW25113 cells with pEvol-*Nn*SULT1C1 -cysDNCQ, pUltra-sTyr, and a plasmid encoding the thrombin inhibitor. In parallel, we utilized the ΔcysH BW25113 cells lacking the sTyr biosynthetic systems but exogenously fed with 3 mM sTyr (**Fig. 5C** and **S9**). The site-specific sulfation of madanin-1 and chimadanin was further validated using ESI-MS analysis (**Fig. S10**).

**Figure 5:**
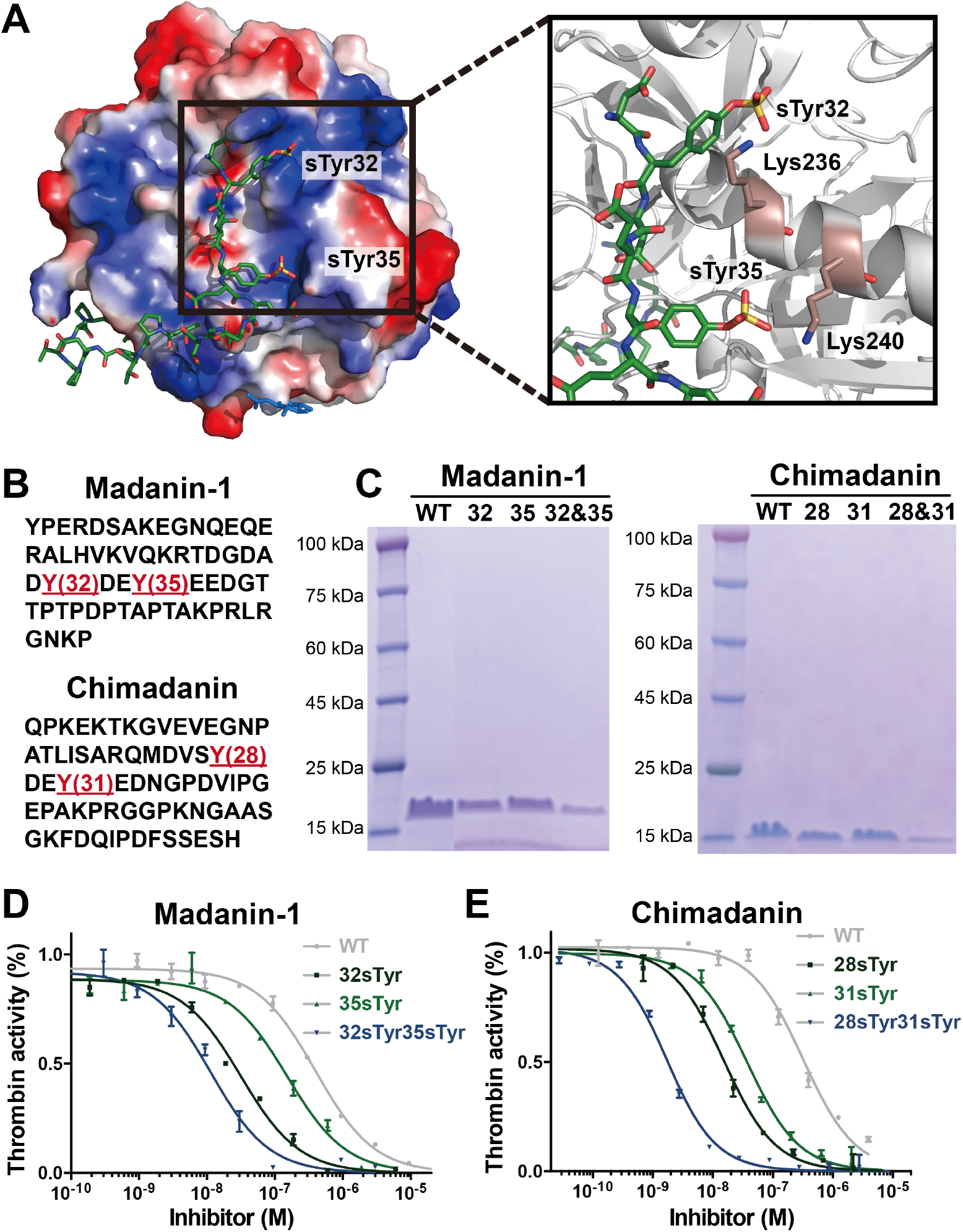
Production of thrombin inhibitors with site-specific sTyr insertion using completely autonomous *E. coli*. **(A)** Madanin-1 with sulfation at Tyr32 and Tyr35 positions binds to exosite II site in thrombin. Surface representation of positive electrostatic potential in blue and negative electrostatic potential in red. **(B)** Amino acid sequences of madanin-1 and chimadanin. Sulfation sites are shown in red. **(C)** SDS-PAGE analysis of thrombin inhibitors with site-specific sTyr insertion expressed in completely autonomous *E. coli* with. **(D-E)** Inhibition of thrombin activity by madanin-1 and chimadanin proteins. Error bars represent standard deviation.

To test the thrombin inhibiting activity of the wild type inhibitors and their sTyr-containing mutants, we performed chromogenic thrombin amidolytic activity assays in the presence of a range of concentrations of each inhibitor. Compared with wildtype madanin-1 (*Ki* = 16.0 ± 0.9 nM), incorporation of a single sTyr at either Tyr32 (*Ki* = 1.3 ± 0.1 nM) or Tyr35 (*Ki* = 6.1 ± 0.6 nM) position significantly enhanced its inhibition of thrombin (**Fig. 5D and S11**). To our delight, madanin-1 mutants sulfated at both Tyr32 and Tyr35 exhibited the highest potency (*Ki* = 0.5 ± 0.1 nM) against thrombin activity (**Fig. 5D and S11**). Following a similar trend, incorporating a single biosynthesized sTyr at either Tyr28 or Tyr31 of chimadanin yields more potent inhibition of thrombin activity (*Ki*= 0.6 ± 0.1 nM and 1.5 ± 0.1 nM, respectively) than achieved with wildtype chimadanin (*Ki* = 12.9 ± 0.1 nM, **Fig. 5E and S11**). Double sulfation of chimadanin at both Tyr28 and Tyr31 further improved its *Ki* to 0.1 nM, consistent with the madanin-1 study. Furthermore, sTyr-containing thrombin inhibitors prepared using cells with completely autonomous sTyr biosynthetic machinery are more potent than chemically synthesized ones.^25^ This may due to the fact that co-translational folding is more efficient than that achieved via chemical synhesis. These data demonstrate the advantages of producing therapeutic proteins with site-specific sTyr modifications using completely autonomous cells with the ability to biosynthesize and genetically encode the sTyr.

## Discussion

In this research, we have created completely autonomous bacterial and mammalian cells endowed with machinery for both sTyr biosynthesis and site-specific incorporation into proteins. *Nn*SULT1C1 sulfotransferase-mediated biosynthesis of sulfotyrosine from tyrosine and PAPS was discovered using a sequence similarity network (SSN), and the unique specificity of *Nn*SULT1C1 for tyrosine was systematically explored using both bioinformatic and computational methods. Use of *Nn*SULT1C1 and other optimized components allowed us to create both bacterial and mammalian cells capable of autonomously biosynthesizing sTyr and genetically incorporating it into proteins. The resulting cells produce site-specifically sulfated proteins at higher yields than cells exogenously fed with 3-27 mM sTyr. The value of these completely autonomous cells was further demonstrated via their use in the preparation of therapeutic sTyr-containing proteins with enhanced efficacy.

More than 300 ncAAs have been genetically incorporated into proteins, providing powerful tools for investigating protein structures and functions.^1–3,7,77–85^ To date, utilizing these ncAAs in the context of Genetic Code Expansion has required both exogenous feeding and good membrane permeability of chemically-synthesized ncAAs. Cell membranes are poorly permeable to ncAAs with charged, highly hydrophobic, or hydrophilic structures. Thus, intracellular biosynthesis of these ncAAs is likely to significantly expand the utility of Genetic Code Expansion technology. Attempts to engineer cells for autonomous ncAA biosynthesis have frequently been hindered by the scarcity of verified biosynthetic pathways for producing ncAAs at high concentrations. For this reason, biosynthetic pathways for *p*AF, *p*Thr, 5HTP, and DOPA are the only ones that have been applied to bacterial cells for intracellular ncAA biosynthesis from simple carbon sources.^14,16–19^ We expect that the combination of bioinformatics and ncAA screening methods reported in this work can be a general strategy for enlarging the repertoire of biosynthesized ncAA for Genetic Code Expansion. Our study further reports the construction of a completely autonomous mammalian cell line capable of biosynthesizing sTyr and incorporating it into proteins in response to the amber codon. The creation of additional mammalian cells with the endogenous ability to biosynthesize ncAAs and use them for protein synthesis will expand the preparation of therapeutic proteins, as well as allow application of the Genetic Code Expansion technology at the level of whole organisms.

## Acknowledgments

We thank Dr. Xiao Laboratory members for insightful comments. This work was supported by the Cancer Prevention Research Institute of Texas (CPRIT RR170014 to H.X.), NIH (R35-GM133706, R21-CA255894, and R01-AI165079 to H.X.), the Robert A. Welch Foundation (C-1970 to H.X.), US Department of Defense (W81XWH-21-1-0789 to H.X.), the John S. Dunn Foundation Collaborative Research Award (to H.X.), and the Hamill Innovation Award (to H.X.), Center for Theroretical Biological Physics (NSF grant PHY-2019745 to P.G.W.) and D. R. Bullard Welch Chair at Rice University (Grant C-0016 to P.G.W). H.X. is a Cancer Prevention & Research Institute of Texas (CPRIT) scholar in cancer research.

## Methods

### Sequence Similarity Network (SSN)

The SSN was generated by inputting the amino acid sequence of *Rn*SULT1A1 as query sequence at https://efi.igb.illinois.edu/efi-est/. The UniProt database was selected and the e-value was set as 5. The resulting network was finalized by setting the alignment score threshold as 110 and used to generate edges representing pairwise sequence similarities. The representative node network with %ID of 80% was downloaded in the format of xgmml and visualized within Cytoscape.

### Optimized expression of sfGFP-sTyr from sTyr biosynthesis

ΔcysH BW25113 cells, transformed with pUltra-sTyrRS, pET22b-T5-sfGFP151TAG, and pEvol-*Nn*SULT1C1-cysDNCQ, were grown in Luria-Bertani (LB) medium at 37°C. When the OD600 of the cell culture reached 0.6, *Nn*SULT1C1 expression was induced by 15 mg/L *l*-arabinose and grown at 30 °C. After 6 h induction, the cells were diluted 5 times to OD 0.6. Expression of reporter sfGFP and sTyrRS were induced with 1 mM IPTG. Additional *l*-arabinose was also added to maintain its final concentration of 15 mg/L. The control cells transformed with pUltra-sTyrRS, pET22b-T5-sfGFP151TAG and pEvol-empty were grown under the same condition with an indicated concentration of sTyr. After growth at 30 °C for 18 hours, cells were harvested by centrifugation at 4,750 × g for 10 min and used for GFP fluorescence and cell optical density measurements. Proteins were purified on Ni-NTA resin (Qiagen) following the manufacturer’s instructions. The purified protein was used for SDS-PAGE and ESI-MS analysis.

### Predicting the structure of *Nn*SULT1C1 by AlphaFold

The structure of *Nn*SULT1C1 was predicted by AlphaFold2 using GitHub AlphaFold code 2.0. The database including, reduced BFD, PDB70, MGnify, and Uniclust30, was used to filter structural templates. All other settings were set as default. Based on pLDDT, the top structure was output and used in this study.

### Protein-ligand docking

The protein-ligand docking process was performed by Glide v8.1 using Schrödinger software package v2018.4. Glide uses the OPLS3 force field to evaluate the docking procedure. OPLS3 is an enhanced version of the OPLS_2005 all-atom force field to provide a larger coverage of organic functionality. Four protein structures, including 2zvq, 2a3r, 2gwh, and the predicted structure of sulfotransferase, are taken into consideration for docking. The PAPS binding site for the predicted *Nn*SULT1C1 structure is inferred by aligning with the structure of 2a3r. For other structures, we use the original PAP sites in reported co-crystal structures to install PAPS. A short run of protein-ligand energy minimization was performed to remove the steric clashes for each of the complexes. Dockings box was inferred from the position of dopamine in 2a3r. The RMSD is set to 0.5 to sample the distinct conformations. All parameters are set to default SP mode in the Glide software. The maximum output poses for each docking protein was set to 200 and the top 50 poses that ranked by Emodel score were picked out. The best docking pose for each complex was compared using the Glide docking score, an empirical scoring function that approximates the ligand binding free energy in the unit of kcal/mol.

### Characterization of HEK293T-*Nn*SULT1C1 with Confocal Microscopy and Flow Cytometry

To generate HEK293T-*Nn*SULT1C1, HEK293T were transfected with PB-*Nn*SULT1C1 (100 ng) and Piggybac transposases plasmids (20 ng) with Polyjet In Vitro DNA Transfection Reagent (SignaGen Laboratories). 1 μg/mL puromycin was added to culture medium from Day 2 to Day 7 for selecting cells with genomic integration of *Nn*SULT1C1. The puromycin concentration was raised to 3 μg/mL from Day 8 and maintained in the future. HEK293T and HEK293T-*Nn*SULT1C1 cells were cultured in DMEM supplemented with 10% fetal bovine serum (FBS) and 1% penicillin/streptomycin at 37°C and 5% CO_2_. HEK293T and HEK293T-*Nn*SULT1C1 cells were transfected with pAcBacw.tR4-sTyrRS/GFP* with Polyjet In Vitro DNA Transfection Reagent (SignaGen Laboratories) in the presence or absence of the indicated concentration of sTyr. Mediums were changed 12-16 hours after transfection. After 48 hours of the transfection, cells were used for confocal microscopy where nucleus staining was performed by incubating cells with 5 uM DRAQ5 for 10 min. After washed with PBS (pH 7.4) for three times, cells were imaged with Zeiss LSM710 confocal microscopy. The rest of cells were used for flow cytometry analysis with Sony SA3800 Flow Cytometer where a total of 20,000 cells were analyzed for each sample. Data were processed with FlowJo. Reported data is the average measurement of three samples prepared at the same time with the standard deviation.

### Thrombin Activity Assay

N-(p-Tosyl)-GPR-pNA acetate (Cayman Chemicals) was used as a chromogenic substrate to test the amidolytic activity of human α-thrombin (Haematologic Technologies). Purified chi and mad inhibitors were buffer-exchanged to the assay buffer (pH 8) containing 50 mM Tris-HCl, 50 mM NaCl using PD-10 columns. Inhibition assays were performed in the assay buffer with 0.14 nM human α-thrombin, 100 μM substrate, and varying concentrations of inhibitors. After incubation at room temperature for 2 h, the activity of thrombin was monitored by absorption at 405 nm. Inhibition constants (K_i_) were determined based on a Morrison equation within GraphPad Prism. Three independent replicates were prepared for each group.

