## Supporting Information for "Unleashing the Potential of Noncanonical Amino Acid Biosynthesis for Creation of Cells with Site-Specific Tyrosine Sulfation"

### Supplemental Figures

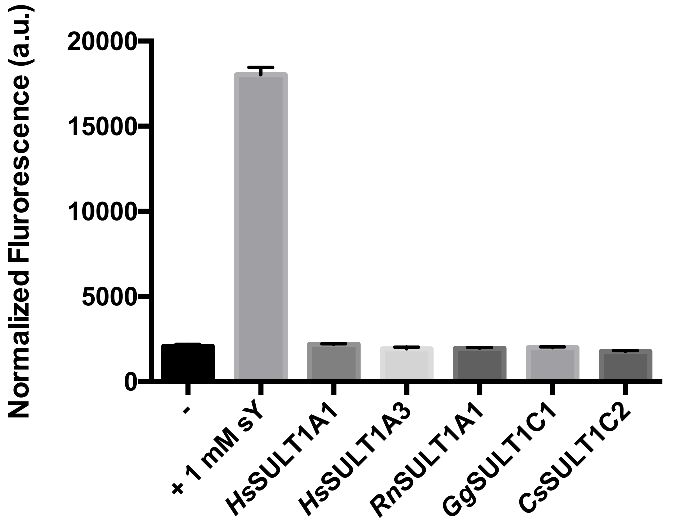

**Figure S1: Screening of reported sulfotransferases with super-folder GFP assay.** Error bar represents strandard deviation.

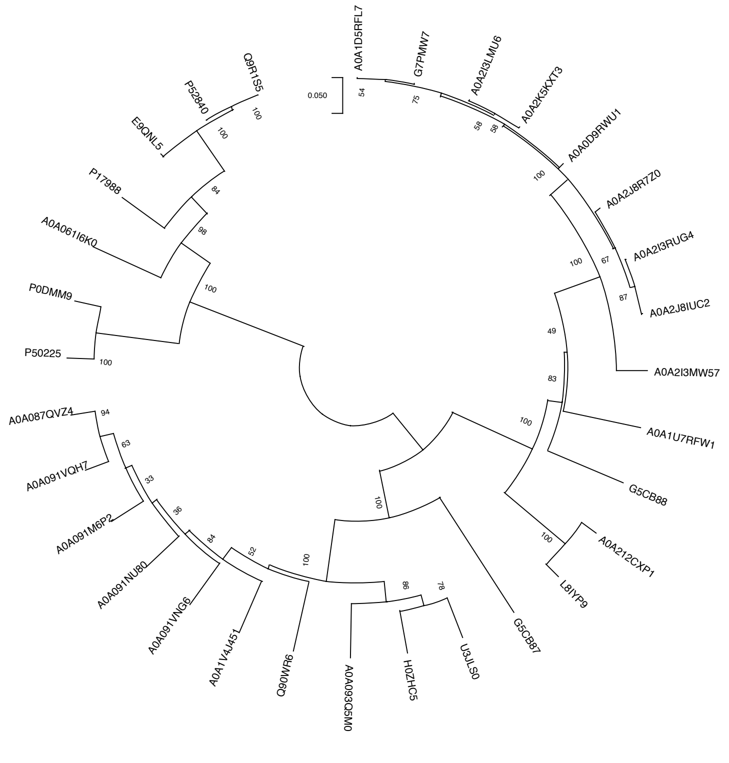

**Figure S2: Phylogenetic relationship of all sulfotransferases tested in Figure. 2D.** Phylogenetic tree was generated in MEGAX software with UPGMA method. A0A091VQH7 (Uniprot ID) was named *Nn*SULT1C1 and used for following experiments.

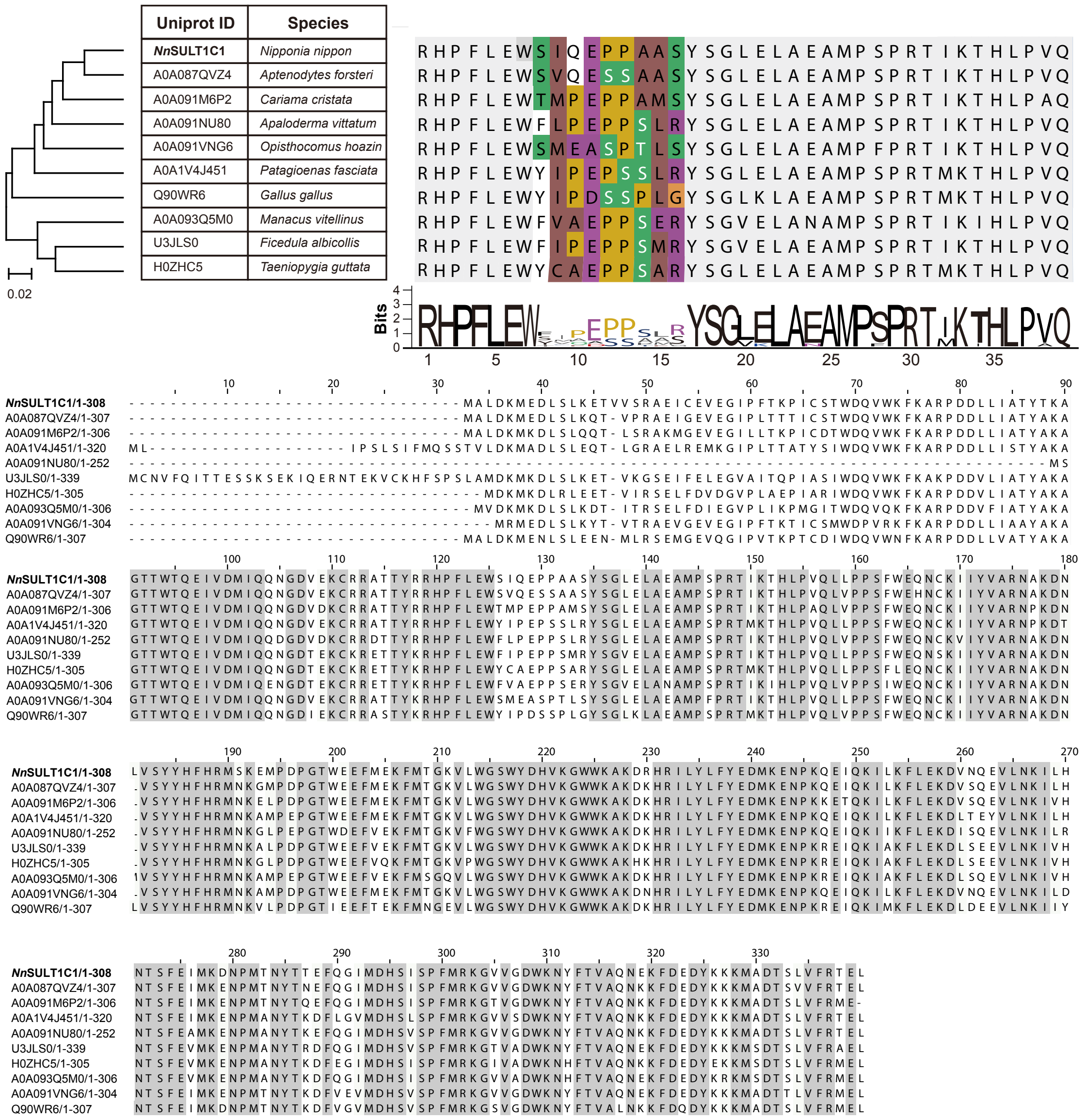

**Figure S3: Phylogenetic tree and multiple sequence alignment analysis of *Nn*SULT1C1 and its 9 relatives.** Phylogenetic tree was constructed by UPGMA method in MEGAX and multiple sequences were aligned by ClustalW method.

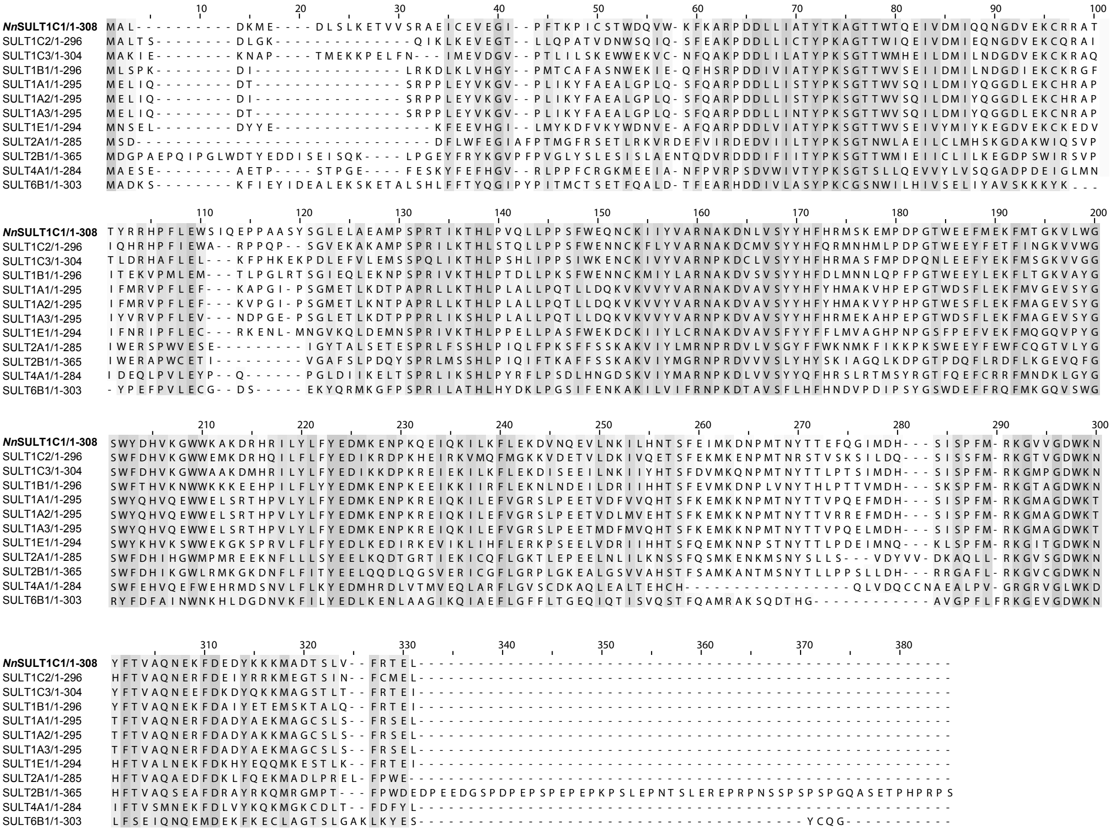

**Figure S4: Sequence alignment of human cytosolic sulfotransferases (hSULTs) and *Nn*SULT1C1.** The highly variable region we found (SIQEPPAAS) in NnSULT1C1 is aligned well with the reported substrate binding loop (colored in cyan) of hSULTs.)

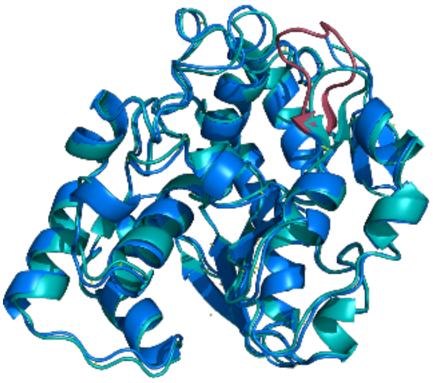

**Figure S5: Superimposition of *Nn*SULT1C1 and 2zvq.** *Nn*SULT1C1 was shown in blue colour and 2zvq (mouse SULT1D1) was shown in cyan colour. The variable region SIQEPPAAS of *Nn*SULT1C1 is shown in red.

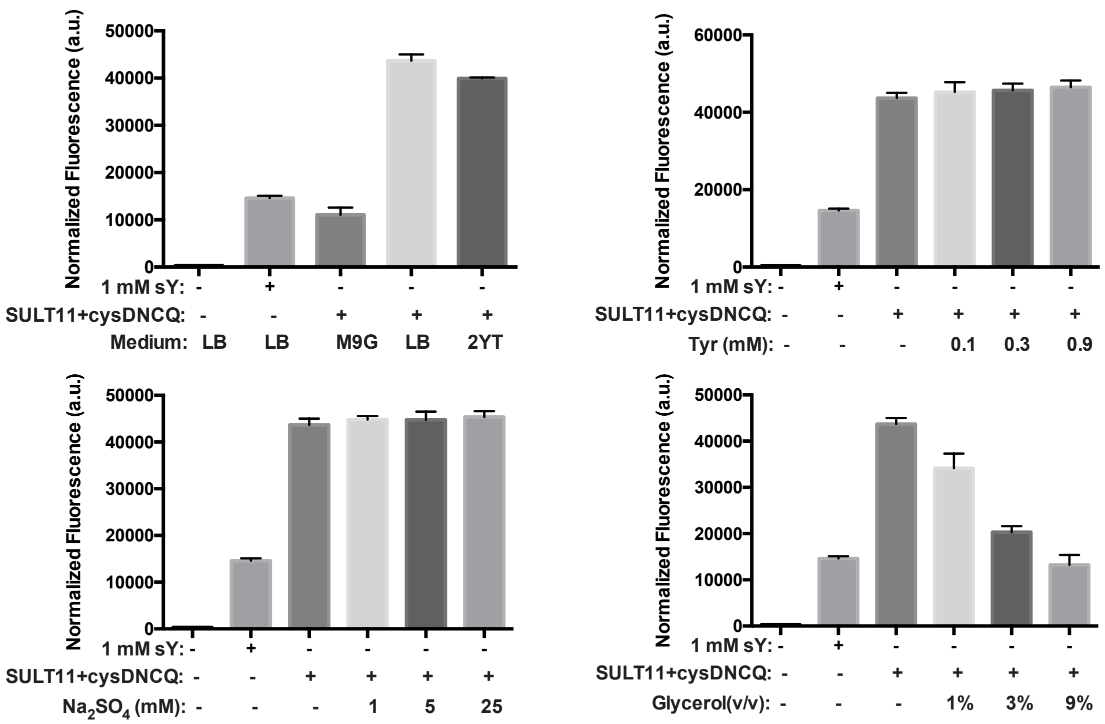

**Figure S6: Expression condition screening for sfGFP-sTyr production.** The influence of expression medium, tyrosine addition, sulfate addition, glycerol addition on production of sfGFP-sTyr in bacterial cells containing sTyr biosyntheis and genetic incorporation machineries was evaluated with green fluorescent protein assay.

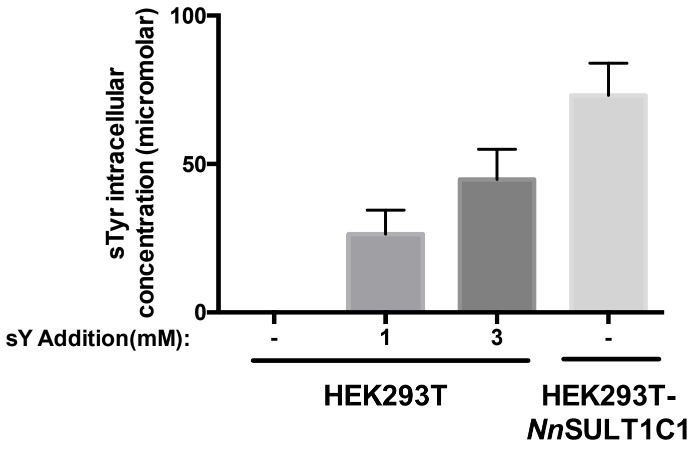

**Figure S7: Cellular concentration of sTyr in HEK293T and HEK293T-*Nn*SULT1C1.** Indicated concentration of sTyr was added to the culture of HEK293T for 2 hour.

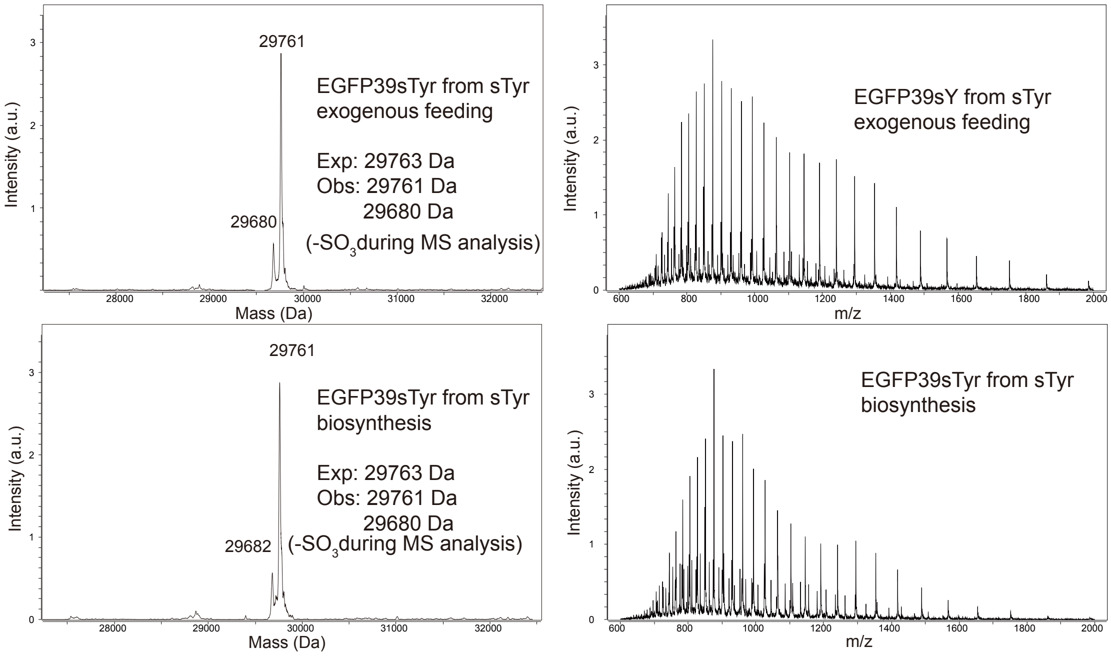

**Figure S8: ESI-MS analysis of EGFP39sTyr from HEK293T cells and HEK293T-*Nn*SULT1C1.** The expected peak was calculated according to monoisotopic mass of EGFP39sY with N-terminal acetylation. Bottom left spectrum is identical to Fig. 4D.

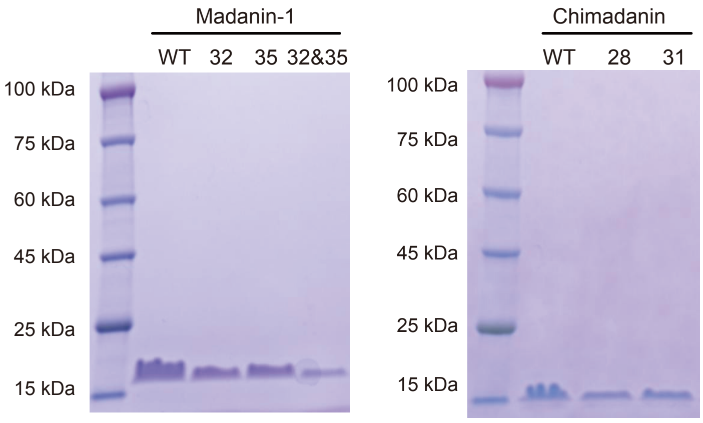

**Figure S9: SDS-PAGE analysis of thrombin inhibitors purified from LB medium.** sTyr-containing inhibitors are expressed in LB medium with external addition of 3 mM sTyr.

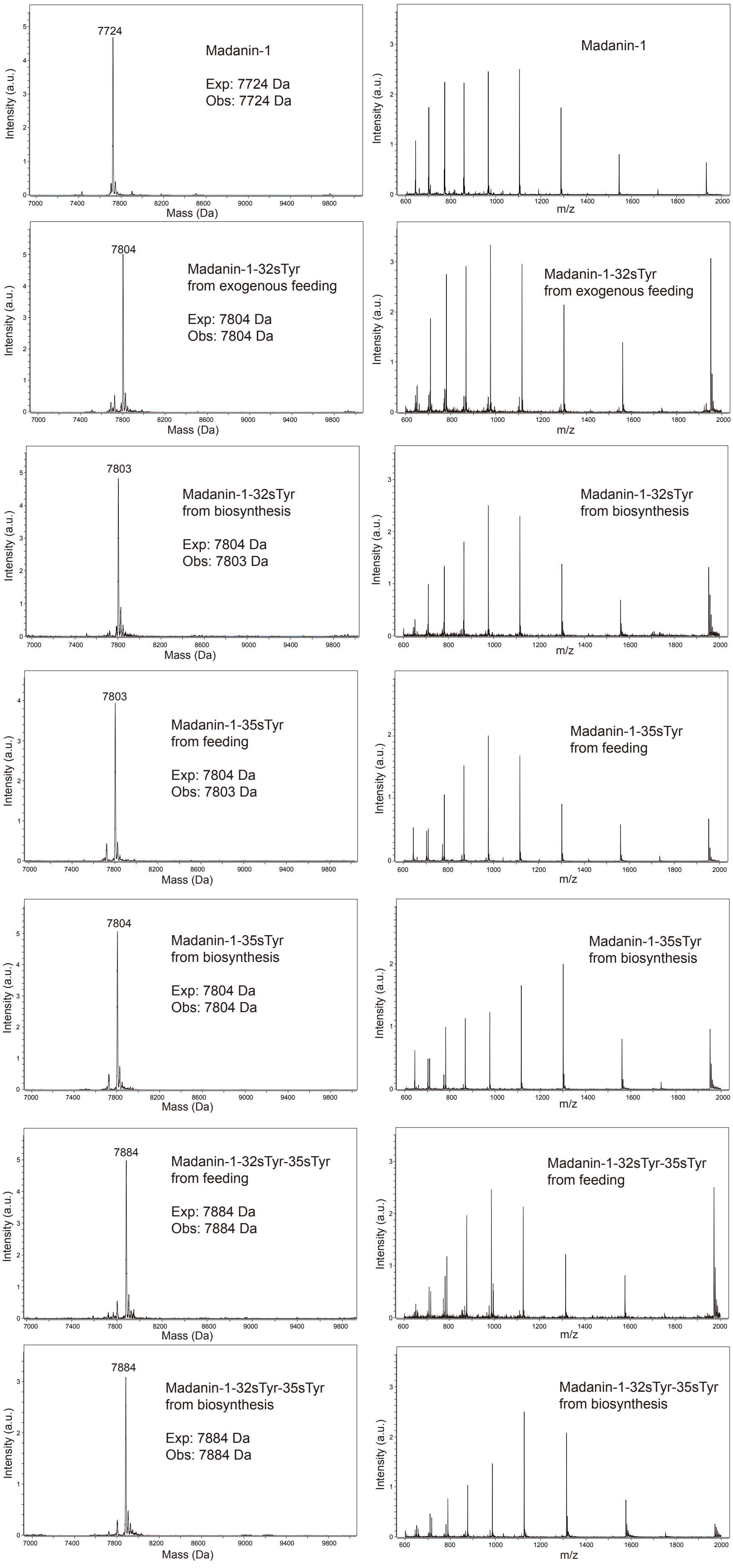
7724
Madanin-1
Exp: 7724 Da
Obs: 7724 Da
Intensity (a.u.)
Mass (Da)
Madanin-1
m/z
7804
Madanin-1-32sTyr
from exogenous feeding
Exp: 7804 Da
Obs: 7804 Da
Madanin-1-32sTyr
from exogenous feeding
7803
Madanin-1-32sTyr
from biosynthesis
Exp: 7804 Da
Obs: 7803 Da
Madanin-1-32sTyr
from biosynthesis
7803
Madanin-1-35sTyr
from feeding
Exp: 7804 Da
Obs: 7803 Da
Madanin-1-35sTyr
from feeding
7804
Madanin-1-35sTyr
from biosynthesis
Exp: 7804 Da
Obs: 7804 Da
Madanin-1-35sTyr
from biosynthesis
7884
Madanin-1-32sTyr-35sTyr
from feeding
Exp: 7884 Da
Obs: 7884 Da
Madanin-1-32sTyr-35sTyr
from feeding
7884
Madanin-1-32sTyr-35sTyr
from biosynthesis
Exp: 7884 Da
Obs: 7884 Da
Madanin-1-32sTyr-35sTyr
from biosynthesis

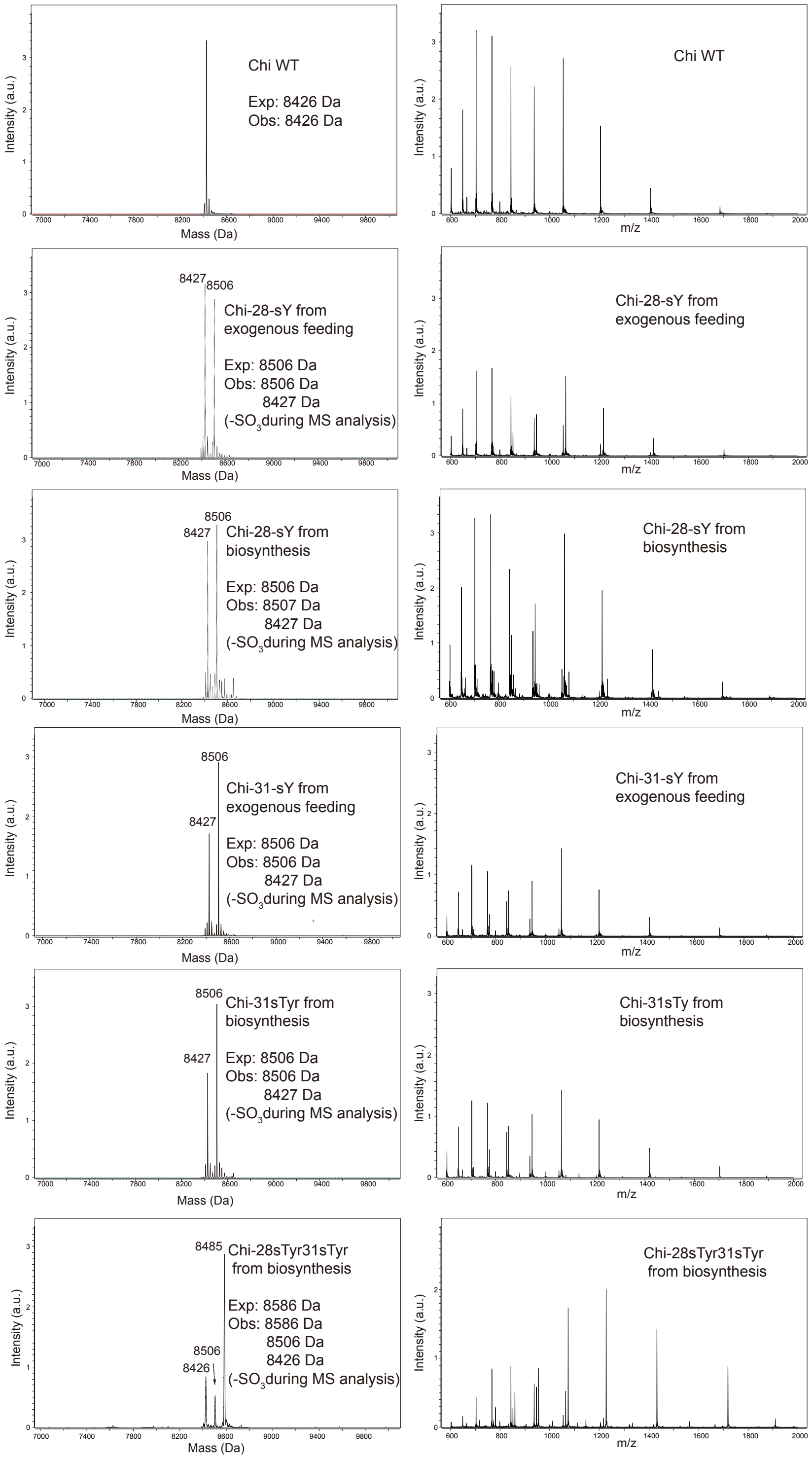

**Figure S10: ESI-MS analysis of all thrombin inhibitors used in this study.**

| Madanin-1 | | | Chimadanin | | |
|---|---|---|---|---|---|
| Form | sTyr source | $K_i$ (nM) | Form | sTyr source | $K_i$ (nM) |
| Wildtype | | 16.0 $\pm$ 0.9 | Wildtype | | 12.9 $\pm$ 1 |
| 32sTyr | Biosynthesis | 1.3 $\pm$ 0.1 | 28sTyr | Biosynthesis | 0.6 $\pm$ 0.1 |
| 35sTyr | | 6.1 $\pm$ 0.6 | 31sTyr | | 1.5 $\pm$ 0.1 |
| 32sTyr35sTyr | | 0.5 $\pm$ 0.1 | 28sTyr31sTyr | | 0.1 $\pm$ 0 |
| 32sTyr | External addition | 1.8 $\pm$ 0.2 | 28sTyr | External addition | 0.5 $\pm$ 0.1 |
| 35sTyr | | 6.0 $\pm$ 0.3 | 31sTyr | | 1.9 $\pm$ 0.2 |
| 32sTyr35sTyr | | 0.3 $\pm$ 0.1 | 28sTyr31sTyr | | |

**Figure S11: Inhibition constants (Ki) of all thrombin inhibitors used in this study.** $K_i$ were calculated based on a tight-binding model, using Morrison equation with Prism.

### Supplemental Experimental Procedures

**Materials**

LB agar and 2YT were obtained from BD Difco™. Isopropyl-β-D-thiogalactoside (IPTG) was ordered from Anatrace. 4-12% Bis-Tris gels for SDS-PAGE were purchased from Invitrogen. Oligonucleotide primers were purchased from Integrated DNA Technologies and Eurofins Genomics (Supplementary Table S1 lists the oligonucleotides used in this report). Plasmid DNA preparation was carried out with the GenCatch™ Plasmid DNA Miniprep Kit and GenCatch™ Advanced Gel Extraction Kit. BugBuster™ Protein Extraction Reagent was obtained from Novagen (Cat. 70584). Pierce™ universal nuclease was purchased from Thermo Scientific (Cat. 88700). $Ni^{2+}$-NTA Agarose was obtained from Qiagen (Cat. 30230). M9-glucose minimal medium contain M9 salt (6.78 g/L $Na_2HPO_4$, 3 g/L $KH_2PO_4$, 1 g/L $NH_4Cl$, 0.5 g/L NaCl), heavy metal solution (1 µg/L $CuSO_4 \cdot 5H_2O$, 4 µg/L $MnCl_2 \cdot 4H_2O$, 4 µg/L $ZnCl_2$, 1.2 µg/L $FeSO_4 \cdot 5H_2O$), 1 mM $MgSO_4$, 0.1 mM $CaCl_2$, 5 µg/mL Thiamine, 300 µM Leucine 4 µM D-Biotin, Glucose (4 g/L).

Unless otherwise mentioned, all solvents and chemicals for synthesis were purchased from Alfa Aesar and Fisher Chemical and used as received without further purification, unless otherwise specified.

**Plasmids Construction**

pUltra-sTyrRS and pAcBacw.tR4-OMeYRS/GFP* were obtained from addgene. pLei-sfGFP134TAG and pEvol-Mj are generous gifts from Dr. Peter Schultz. Piggybac vector and Piggybac transposase plasmids are kind gifts from Dr. Caleb Bashor.

Sulfotransferase containing pBad plasmids for initial screening were constructed by Gibbson Assembly of pBad Vector amplified from pBad-HER2-ScFv with Da343&344 and sulfotransferase fragment amplified from synthetic DNA with respective primers shown in Table S1. pEvol-*Nn*SULT1C1 was acquired by Gibbson Assembly of pEvol-Mj vector amplified by Da443&444 and *Nn*SULT1C1 amplified from pBad-*Nn*SULT1C1with Da441&442. To generate pEvol-cysDNC, cysDNC cassette was amplified from *E.coli* genome with Da446 and

Da447 and inserted into pEvol vector amplified from pEvol-Mj with Da443&445. To generate pEvol-cysDNCQ, cysDNC cassette and cysQ cassette amplified seperately from *E.coli* genome with Da446&448 and Da449&450, respectively, were overlapped and inserted into pEvol vector amplified from from pEvol-Mj with Da443&445. pEvol-*Nn*SULT1C1-cysDNC(Q) were generated by inserting *Nn*SULT1C1 fragment amplified from pBad-*Nn*SULT1C1 into the vector of pEvol-cysDNC(Q) amplified with Da444&454.

To generate the plasmid for sTyr genetic incorporation in mammalian cells, OMeYRS of pAcBacw.tR4-OMeYRS/GFP* was substituted with sTyr-selective EcTyrRS mutant (L71V, D182G, L186M), which was achieved by Gibbson Assembly of synthetic sTyrRS amplified with Da462&463 and pAcBacw.tR4-OMeYRS/GFP* vector digested by Xho1 and Nhe1. To integrate *Nn*SULT1C1 into genome of HEK293T, PB-*Nn*SULT1C1 was constructed by assembling synthetic *Nn*SULT1C1 amplified by Da655&656 and Piggybac vector digested with BsrG1.

pET22b-T5-chi was constructed by Gibbson Assembly of synthetic chimadanin coding sequence amplified with Da556&557 and pET22b-T5-sfGFP151TAG vector digested by Hind3 and Nde1. pET22b-T5-chi-28TAG and pET22b-T5-chi-31TAG were made according to the protocol on NEBaseChanger with Da664&665 and Da584&585, respectively, using pET22b-T5-chi as their template. pET22b-T5-chi-28TAG31TAG were made by following the same protocol with Da666&Da664 with pET22b-T5-chi-31TAG as a template.

pET22b-T5-mad was constructed by Gibbson Assembly of synthetic madanin-2 coding sequence amplified with Da558&559 and pET22b-T5-sfGFP151TAG vector digested by Hind3 and Nde1. pET22b-T5-mad-32TAG and pET22b-T5-mad-35TAG were made according to the protocol on NEBaseChanger with Da586&587 and Da661&662 with pET22b-T5-mad as a template. pET22b-T5-mad-32TAG35TAG were made by following the same protocol with Da663&Da661 using pET22b-T5-mad-32TAG as a template.

To explore whether sulfotransferase with similar structure with *Nn*SULT1C1 could catalyze tyrosine sulfation, pEvol-mSULT1D1/hSULT1C2-cysDNCQ was constructed by assembling synthetic sulfotransferases amplified by Da832&833/ Da838&839 with pEvol-*Nn*SULT1C1-cysDNCQ vector amplified with Da802&803.

To test the importance of *Nn*SULT1C1 loop (SIQEPPAASY) and residues (T30, I33, W93, E161) involved in substrate binding, their corresponding mutants was obtained by assembly of pEvol-*Nn*SULT1C1-cysDNCQ fragments amplified with Da858&859, Da871&872, Da873&874, Da875&876, Da877&878.

| Oligonucleotide | Sequence (5'-3') | Note |
|---|---|---|
| Da343 | cataaaatcacctcaacctctagatacc | |
| Da344 | taagtcgaccgatgcccttgag | |
| Da326 | gaattcattaaagaggagaaattacatATGGAATTGATTCAAGATACGAGCCGCC | HsSULT1A1 |
| Da327 | gctctcaagggcatcggtcgacttaTCACAGCTCACTACGAAAGCTAAGCGAG | HsSULT1A1 |
| Da328 | gaattcattaaagaggagaaattacatATGGAGTTCTCTCGCCCTCCACTTGTGC | HsSULT1A3 |
| Da329 | gctctcaagggcatcggtcgacttaTCACAATTCGCAACGAAACTTGAAATCG | HsSULT1A3 |
| Da330 | gaattcattaaagaggagaaattacatATGGAACTTATCCAGGATACCTCCCGTC | RnSULT1A1 |
| Da331 | gctctcaagggcatcggtcgacttaTCAAAGCTCCGAGCGAAACGACAGTGAAC | RnSULT1A1 |
| Da332 | gaattcattaaagaggagaaattacatATGGCTCTGGACAAGATGGAAAACTTG | GgSULT1C1 |

| | | |
|---|---|---|
| Da333 | gctctcaagggcatcggtcgacttaTCAAAGTTCCATACGGAAAACCAAGCTAG | GgSULT1 C1 |
| Da345 | gtatctagaggttgaggtgattttATGGCCTTAACATCTGACCTTGGTAAG | O00338 |
| Da346 | gctctcaagggcatcggtcgacttaTCAGAGTTCCATGCAGAAGTTGATGC | O00338 |
| Da347 | gtatctagaggttgaggtgattttATGGCCTTAACTTCAGAGTTAGGGAAAC | A0A1D5RF L7 |
| Da348 | gctctcaagggcatcggtcgacttaTCACAGCTCCATACAGAAGTTAATACTTGTG | A0A1D5RF L7 |
| Da349 | gtatctagaggttgaggtgattttATGGCTCAGGTTCCTGAATTATCGAAACCG | A0A1U7RF W1 |
| Da350 | gctctcaagggcatcggtcgacttaTCACAGTTTCATACAAAAGTTAATCGAAGTGC | A0A1U7RF W1 |
| Da351 | gtatctagaggttgaggtgattttATGCTTTTAATCAGCACATATGCAAAAGCG | A0A1V4J4 51 |
| Da352 | gctctcaagggcatcggtcgacttaTCATTACAGTTCTGTACGGAAAACAAGTGAAG | A0A1V4J4 51 |
| Da353 | gtatctagaggttgaggtgattttATGATTGAGCAAAACGGGGACGTG | A0A2I3LM U6 |
| Da354 | gctctcaagggcatcggtcgacttaTCACAGTTCCATACAAAAATTGATCGAGGTC | A0A2I3LM U6 |
| Da355 | gtatctagaggttgaggtgattttATGGCATTAACATCTGAGCTTGGTAAGC | A0A2I3MW 57 |
| Da356 | gctctcaagggcatcggtcgacttaTCACAGTTCCATGCAGAAGTTAATACTTGTCC | A0A2I3MW 57 |
| Da357 | gtatctagaggttgaggtgattttATGGATATGATTGAGCAGAACGGCG | A0A2I3RU G4 |
| Da358 | gctctcaagggcatcggtcgacttaTCACAGCTCCATGCAGAAATTAATTGAAGTACC | A0A2I3RU G4 |
| Da359 | gtatctagaggttgaggtgattttATGGCGTTAACCTCGGACTTGG | A0A2J8IU C2 |
| Da360 | gctctcaagggcatcggtcgacttaTCACAGTTCCATACAAAAGTTAATAGCCGTCC | A0A2J8IU C2 |
| Da361 | gtatctagaggttgaggtgattttATGGCTCTGACTTCTGAACTGGGGAAAC | A0A2J8R7 Z0 |
| Da362 | gctctcaagggcatcggtcgacttaTCACAGCTCCATACAAAAATTAATTGCAGTGC | A0A2J8R7 Z0 |
| Da363 | gtatctagaggttgaggtgattttATGGCATTAACAAGCGAGTTGGGGAAG | A0A2K5KX T3 |
| Da364 | gctctcaagggcatcggtcgacttaTCACAGCTCCATGCAAAAGTTAATGCTTGTG | A0A2K5KX T3 |
| Da365 | gtatctagaggttgaggtgattttATGGCTCTCGACAAAATGGAGGAC | A0A091VQ H7 |
| Da366 | gctctcaagggcatcggtcgacttaTCACAGCTCAGTGCGAAAGACCAAC | A0A091VQ H7 |
| Da367 | gtatctagaggttgaggtgattttATGGCCCTTATCACTGCAGGTACTC | A0A212CX P1 |
| Da368 | gctctcaagggcatcggtcgacttaTCACAGTTCTGTGCAAAAATTGATGCTGGTAC | A0A212CX P1 |
| Da369 | gtatctagaggttgaggtgattttAACATGGAGCTTATCAAAGACATTTCGCG | A0A061I6K 0 |
| Da370 | gctctcaagggcatcggtcgacttaTCACAGGTTGCACCGAAATTTGAGG | A0A061I6K 0 |
| Da371 | gtatctagaggttgaggtgattttATGGCGCAAAATCCAAGCAACATG | E9QNL5 |
| Da372 | gctctcaagggcatcggtcgacttaTCAAATTTGACAACGAAATGTGAAATCGCAACCG | E9QNL5 |
| Da373 | gtatctagaggttgaggtgattttATGGAACCACTGCGGAAGCC | P52840 |
| Da374 | gctctcaagggcatcggtcgacttaTCAAATTTGGCACCGAAAAGTGAAGTCG | P52840 |
| Da375 | gtatctagaggttgaggtgattttATGGCCCAGAACCCATCTAACATG | Q9R1S5 |
| Da376 | gctctcaagggcatcggtcgacttaTCAAATTTGACACCGGAATGTGAAGTCAC | Q9R1S5 |
| Da377 | gtatctagaggttgaggtgattttATGGCGGGCGAAGATCACAC | G5CB87 |
| Da378 | gctctcaagggcatcggtcgacttaTCAAATTTCAGTGCGGAATGTAAGGGTAC | G5CB87 |
| Da379 | gtatctagaggttgaggtgattttATGCGGAAACCTGAGCTGGAG | G5CB88 |
| Da380 | gctctcaagggcatcggtcgacttaTCACAGTTCGAGGCAAAACCGG | G5CB88 |
| Da381 | gtatctagaggttgaggtgattttATGGCCCTCACTAGCGAACTTG | G7PMW7 |
| Da382 | gctctcaagggcatcggtcgacttaTCACAGTTCCATACAGAAGTTAATGGACGTG | G7PMW7 |
| Da383 | gtatctagaggttgaggtgattttATGGCTTTAACTACCGCGGGTAC | L8IYP9 |
| Da384 | gctctcaagggcatcggtcgacttaTCACAGTTCGGTGCAGAAGTTAATACTTGTC | L8IYP9 |
| Da425 | gtatctagaggttgaggtgattttATGGCTTTGGATAAGATGGAAGACCTCTC | A0A087QV Z4 |

| | | |
|---|---|---|
| Da426 | gctctcaagggcatcggtcgacttaTCAGAGTTCTGTGCGGAAGACTACG | A0A087QVZ4 |
| Da427 | gtatctagaggttgaggtgattttATGGCGCTGGACAAAATGAAGGAC | A0A091M6P2 |
| Da428 | gctctcaagggcatcggtcgacttaTCACTCCATACGAAAGACCAGAGATGTG | A0A091M6P2 |
| Da429 | gtatctagaggttgaggtgattttATGCGTATGGAAGATCTTTCGCTGAAATAC | A0A091VNG6 |
| Da430 | gctctcaagggcatcggtcgacttaTCACAATTCCATGCGGAAGACTAACG | A0A091VNG6 |
| Da433 | gtatctagaggttgaggtgattttATGTGCAATGTGTTCCAGATCACTACCG | U3JLS0 |
| Da434 | gctctcaagggcatcggtcgacttaTCACAACTCTGCCCGAAACACC | U3JLS0 |
| Da435 | gtatctagaggttgaggtgattttATGGTAGACAAAATGAAAGACCTCTCACTC | A0A093Q5M0 |
| Da436 | gctctcaagggcatcggtcgacttaTCATAATTCCATGCGGAAAACCAACGAG | A0A093Q5M0 |
| Da437 | gtatctagaggttgaggtgattttATGCTGGCCATGGACAAGATGAAAG | H0ZHC5 |
| Da438 | gctctcaagggcatcggtcgacttaTCACAATTCCATACGGAAGACGAGGC | H0ZHC5 |
| Da439 | gtatctagaggttgaggtgattttATGTCTGGCACTACATGGACTCAGG | A0A091NU80 |
| Da440 | gctctcaagggcatcggtcgacttaTCAAAGTTCCGTCCGGAAAACAAGTG | A0A091NU80 |
| Da443 | gcatgctcgagcagctcag | |
| Da444 | agatctaattcctcctgttagcccaaaaaaacg | |
| Da441 | tgggctaacaggaggaattagatctATGGCTCTCGACAAAATGGAGGAC | |
| Da442 | cgaccctgagctgctcgagcatgcaGAAGACAGTCATAAGTGCGGCGA | |
| Da445 | atgggattcctcaaagcgtaaacaacgtataac | |
| Da446 | gtttacgctttgaggaatcccatATGGATCAAATACGACTTACTCACCTGCG | |
| Da447 | cgaccctgagctgctcgagcatgcTCAGGATCTGATAATATCGTTCTGTCTCAACAG | |
| Da448 | TCAGGATCTGATAATATCGTTCTGTCTCAACAG | |
| Da449 | gacagaacgatattatcagatcctgaTAAGTTAACACCGCTCACAGAGACGAG | |
| Da450 | cgaccctgagctgctcgagcatgcTTAGTAAATAGACACTCTGAACCCCGGATTC | |
| Da454 | gtcgaccatcatcatcatcatcattgagtttaaac | |
| Da462 | TCACTATAGGGAGACCCAAGCTGGCTAGCGCCACCATGGCGAGTTCCAATCTGATTAAGC | |
| Da463 | GAGTTAAAGTCGACTTAACGCGTTGAATTCTTATACAGGTCCTTTCCAGCAAATGAGAC | |
| Da556 | attcattaaagaggagaaattacatATGCAGCCTAAGGAAAAAACAAAAGGTGTAG | |
| Da557 | gagtccaagctcagctaattaagcttATGAGATTCAGAACTGAAGTCAGGAATCTGATC | |
| Da558 | attcattaaagaggagaaattacatATGTATCCAGAACGGGACTCCG | |
| Da559 | gagtccaagctcagctaattaagcttCGGTTTGTTGCCCCGAAGG | |
| Da584 | TTACGACGAGtagGAGGACAACGC | |
| Da585 | CTAACGTCCATCTGACGGGCG | |
| Da586 | GGACGCGGATtagGATGAATATGAGGAAG | |
| Da587 | CCGTCGGTACGTTTCTGCACC | |
| Da655 | GGCAAAGAATTGCAAGTTTGTACAAAAAAGCAGGCTGCCACCATGGCGCTTGACAAAATGGAAGAC | |
| Da656 | GCCTGCACCTGAGGATCACCACTTTGTACAAGAAAGCTGGGTTTAaagctctgtccgaaagacgagagaag | |
| Da661 | TCCGCGTCCCCGTCGGTA | |
| Da662 | TTACGATGAAtagGAGGAAGACGGGACGAC | |
| Da663 | TTAGGATGAAtagGAGGAAGACGGGACGAC | |
| Da664 | ATCTGACGGGCGGAAATGAGCG | |
| Da665 | GGACGTTAGTtagGACGAGTACGAGGACAACG | |
| Da666 | GGACGTTAGTtagGACGAGTAGGAGGACAACG | |
| Da858 | CGGCACCCGTTCCTCGAATGGTCTGGGCTTGAATTAGCGGAGGC | |
| Da859 | CCATTCGAGGAACGGGTGCC | |
| Da871 | GAAGTCGAAGGTATCCCGTTCGCGAAGCCTATTTGTAGTACGTGGGATCAAGTG | |
| Da872 | GAACGGGATACCTTCGACTTCGCAG | |
| Da873 | GAAGGTATCCCGTTCACTAAGCCTGCGTGTAGTACGTGGGATCAAGTGTGGAAATTC | |
| Da874 | AGGCTTAGTGAACGGGATACCTTCG | |
| Da875 | GCCGGCACCCGTTCCTCGAAGCGTCAATCCAGGAGCCACCGGCT | |
| Da876 | TTCGAGGAACGGGTGCCGGC | |
| Da877 | CTATCACTTTCACCGCATGAGCAAAGCGATGCCAGATCCTGGGACCTGG | |
| Da878 | TTTGCTCATGCGGTGAAAGTGATAGTAAC | |

**Expression and Purification of Proteins.**

*E. coli* BL21(DE3) cells, transformed with pUltra-sTyrRS, pLei-sfGFP134TAG, and pBad-Empty/pBad-*Hs*SULT1A1/pBad-*Hs*SULT1A3/pBad-*Rn*SULT1A1/pBad-*Gg*SULT1C1/pBad-*Cs*SULT1C2, were grown in 2YT medium at 37°C. The protein expression was carried out in Luria-Bertani (LB) medium with or without 1 mM sTyr addition. When the OD600 of the cell culture reached 0.6, protein expression was induced by the addition of IPTG and *l*-arabinose to a final concentration of 1 mM and 0.2%, respectively. After growth overnight at 30 °C. Cells were harvested by centrifugation at 4,750 × g for 10 min and used for GFP fluorescence and cell optical density measurements. (Fig.S1)

BW25113, ΔtrpE BW25113, ΔtyrA BW25113, ΔackA BW25113, ΔptsH BW25113, ΔcysH BW25113 cells transformed with pUltra-sTyrRS, pET22b-T5-sfGFP151TAG, and pEvol-*Nn*SULT1C1/pEvol-Empty, were grown in 2YT medium at 37°C. The protein expression was carried out in Luria-Bertani (LB) medium with or without 1 mM sTyr addition. When the OD600 of the cell culture reached 0.6, protein expression was induced by the addition of IPTG and *l*-arabinose to a final concentration of 1 mM and 0.2%, respectively. After growth overnight at 30 °C. Cells were harvested by centrifugation at 4,750 × g for 10 min and used for GFP fluorescence and cell optical density measurements. (Figure. 3b)

ΔcysH BW25113, transformed with pUltra-sTyrRS, pET22b-T5-sfGFP151TAG, and pEvol-Empty/pEvol-*Nn*SULT1C1/ pEvol- *Nn*SULT1C1-cysDNC/pEvol- *Nn*SULT1C1-cysDNCQ, were grown in 2YT medium at 37°C. The protein expression was carried out in Luria-Bertani (LB) medium with or 1 mM sTyr addition. When the OD600 of the cell culture reached 0.6, protein expression was induced by the addition of IPTG and l-arabinose to a final concentration of 1 mM and 0.2%, respectively. After growth overnight at 30 °C. cells were harvested by centrifugation at 4,750 × g for 10 min and used for GFP fluorescence and cell optical density measurements. (Figure. 3c)

ΔcysH BW25113 cells, transformed with pUltra-sTyrRS, pET22b-T5-sfGFP151TAG, and pEvol-Empty/pEvol-*Nn*SULT1C1-cysDNCQ, were grown in 2YT medium at 37°C. The protein expression was carried out in Luria-Bertani (LB) medium. When the OD600 of the cell culture reached 0.6, *Nn*SULT1C1 expression was induced by 15 mg/L *l*-arabinose and grown at 30°C for 6 h. Then the cells were diluted 5 times to OD 0.6. Expression of reporter sfGFP and sTyrRS were induced with 1 mM IPTG and indicated concentration of sTyr was added at same time. Additional *l*-arabinose was also added to maintain its final concentration of 15 mg/L. After growth overnight at 30 °C for 18 hours, cells were harvested by centrifugation at 4,750 × g for 10 min and used for GFP fluorescence and cell optical density measurements. (Figure. 3d)

ΔcysH BW25113 cells, transformed with pUltra-sTyrRS, pET22b-T5-sfGFP151TAG, and pEvol-Empty/pEvol-*Nn*SULT1C1-cysDNCQ, were grown in 2YT medium at 37°C. The protein expression was carried out in Luria-Bertani (LB) medium. When the OD600 of the cell culture reached 0.6, *Nn*SULT1C1 expression was induced by indicated concentration of *l*-arabinose and grown at 30°C for 6 h. Then the cells were diluted 5 times to OD 0.6. Expression of reporter sfGFP and sTyrRS were induced with 1 mM IPTG and indicated concentration of sTyr was added at same time. Additional *l*-arabinose was also added to maintain its indicated concentration. After growth at 30 °C for 18 hours, cells were harvested by centrifugation at 4,750 × g for 10 min and used for

GFP fluorescence and cell optical density measurements. (Figure. 3e) Proteins were purified on Ni-NTA resin (Qiagen) following the manufacturer's instructions. The purified protein was used for SDS-PAGE and ESI-MS analysis. (Figure. 3f-h)

To express wildtype thrombin inhibitors, BL21(DE3) cells ransformed with either pET22b-T5-chi or pET22b-T5-mad were grown in 2YT medium at 37°C. The protein expression was carried out in Luria-Bertani (LB) medium. When the OD600 of the cell culture reached 0.6, protein expression was induced by the addition of 0.4 mM IPTG. After growth overnight at 18 °C for 18 hours, cells were harvested by centrifugation at 4,750 × g for 10 min. Proteins were purified on Ni-NTA resin (Qiagen) following the manufacturer's instructions. The purified protein was used for SDS-PAGE and ESI-MS analysis.

To express thrombin inhibitors containing sTyr, ΔcysH BW25113 cells, transformed with pUltra-sTyrRS, pET22b-T5-inhibitor-X-TAG, and pEvol-*Nn*SULT1C1-cysDNCQ, were grown in 2YT medium at 37°C. In the control group, ΔcysH BW25113 cells were transformed with pUltra-sTyrRS, pET22b-T5-inhibitor-X-TAG, and pEvol-Empty. When the OD600 of the cell culture reached 0.6, *Nn*SULT1C1 expression was induced by 15 mg/L concentration of *l*-arabinose and grown at 30°C for 6 h. Then the cells were diluted 5 times to OD 0.6. Expression of reporter inhibitor and sTyrRS were induced with 1 mM IPTG and 3 mM sTyr was added to only control cells at same time. Additional *l*-arabinose was also added to maintain its final concentration of 15 mg/L. After growth overnight at 18 °C for 18 hours, cells were harvested by centrifugation at 4,750 × g for 10 min and used for GFP fluorescence and cell optical density measurements. (Figure. 5c) Proteins were purified on Ni-NTA resin (Qiagen) following the manufacturer's instructions. The purified protein was used for SDS-PAGE and ESI-MS analysis.

To test the importance of the variable loop and residues in binding pockets, ΔcysH BW25113 cells, transformed with pUltra-sTyrRS, pET22b-T5-sfGFP151TAG, and pEvol-*Nn*SULT1C1-cysDNCQ with indicated sequence mutations were grown in 2YT medium at 37°C. The protein expression was carried out in Luria-Bertani (LB) medium. When the OD600 of the cell culture reached 0.6, *Nn*SULT1C1 expression was induced by indicated concentration of *l*-arabinose and grown at 30°C for 6 h. Then the cells were diluted 5 times to OD 0.6. Expression of reporter sfGFP and sTyrRS were induced with 1 mM IPTG and indicated concentration of sTyr was added at same time. Additional *l*-arabinose was also added to maintain its final concentration of 15 mg/L. After growth overnight at 30 °C for 18 hours, cells were harvested by centrifugation at 4,750 × g for 10 min and used for GFP fluorescence and cell optical density measurements. (Figure. 2b-c, f)

**Expression and Fluorescence Measurement of sfGFP**

After sfGFP expression with the same method described above, 0.5 mL cells were harvested by centrifugation at 4,750 × g for 10 min and then suspended with 0.5 ml PBS (pH 7.4). Fluorescence of cells was measured using excitation/emission wavelengths of 395/509 nm. Optical Density at 600 nm was also recorded. The sfGFP fluorescence/OD600 was used as the normalized fluorescence. The error bars in Fig. 3A-C represent for the standard deviation of 3 independent protein expression trials.

***E. coli* Intracellular sTyr Concentration Measurement**

Cells were harvested by centrifugation at 4,750 × g for 10 min and washed with PBS 7.4 for three times. The cell pellets were re-suspended in 300 μL of bugbuster lysis buffer : toluene (80: 20) solution and shaken at 30 ℃

for 1 h. The resulting lysate was centrifuged at 21000 g for 30 min at 4 ℃. 200 µl supernatant was transferred to a new tube and re-centrifuged at 21000 g for 2 h. 50 µl supernatant from the top was then analyzed using the LC-MS. An Agilent 1260 Infinity II LC System coupled with Single Quadrupole ESI-MS System was used for analysis of all samples. 2.5 µL of sample was injected into C18 column. To measure the sTyr ions, ions detected were set to selected ion monitoring (SIM) mode (262 m/z) to detect positive ions of sTyr. Standards of 1 µM, 25 µM, 50 µM, 100 µM, 200 µM, and 400 µM of purchased sTyr (Bachem) were also prepared and injected at a volume of 5 µL by the same method. Using LC-MS data, a linear standard curve was generated based on peak areas corresponding to sTyr ions and the concentration of sTyr in standard samples. The standard curve was then used to calculate the concentrations of sTyr from different cell lysates. The intracellular concentration of sTyr in cells was calculated based on the following equation.

$$[\boldsymbol{sTyr\ intracellular}] = \frac{\boldsymbol{sTyr\ concentration\ in\ lysate\ \times volumn\ of\ lysate}}{\boldsymbol{total\ cell\ numbers\ \times\ E.coli\ cell\ volumn}}$$

Total cell numbers were calculated with the approximate values: $8 \times 10^8$ cells per OD600. 0.6 fL was used as an average *E. coli* cell volume.

**Exploration on evolutionary relationship of *Nn*SULT1C1**

The rooted phylogenetic tree was inferred using the UPGMA method in MEGA X software. The UPGMA algorithm constructs the tree that reflects the genetic distance between protein sequences present in a pairwise similarity matrix. The tree is drawn to scale, with branch lengths in the same units as those of the evolutionary distances used to infer the phylogenetic tree. The evolutionary distances were computed using the Poisson correction method and are in the units of the number of amino acid substitutions per site. 10 sequences from bottom branch were used for Multiple sequence alignment (MSA) on https://www.ebi.ac.uk/Tools/msa/ with Maftt method. The alignment result was visulized in Jalview software. The sequence consensus was analyzed on https://weblogo.berkeley.edu.

**Protein Purification from Mammalian Cells**

To confirm the genetic incorporation of sTyr from either biosynthesis or external addition, HEK293T and HEK293T-*Nn*SULT1C1 cells were transfected with pAcBacw.tR4-sTyrRS/GFP* with Polyjet In Vitro DNA Transfection Reagent (SignaGen Laboratories) in the presence or absence of 3 mM sTyr addition. Mediums were changed at 12-16 hour after transfection. After 48 hours of transfection, cells were harvested with trypsin and subsequently washed by DPBS for 3 times. Cells were lysed using the Mammalian Cell PE LB reagent (G-Bioscience) according to its manual. The cell lysates were centrifuged at 15,000 rpm for 10 minutes. The protein was purified from the supernatant using Ni-NTA resin (Qiagen) following the manufacturer's instruction. The purified protein was used for ESI-MS analysis.

**Mammalian Cell sTyr Concentration Measurement**

HEK293T and HEK293T-*Nn*SULT1C1 were detached from plate with trypsin and washed with DPBS for 3 times. The number of cells was counted by hemocytometer. Cells were resuspended in 0.5 mL methanol-water (2:3) and lyzed by six freeze-thaw cycles. The resulting cell lysates were centrifuged at 21000 × g for 1h at 4℃. The resuting supernantants were injected to LC-MS for the quantification of sTyr with selected ion monitoring (SIM) mode. An Agilent 1260 Infinity II LC System coupled with Single Quadrupole ESI-MS System was used for the analysis of all samples. 10 µL of sample was injected into C18 column. To measure the sTyr ions, ions detected were set to selected ion monitoring (SIM) mode (262 m/z) to detect positive ions of sTyr. Standards of 1 µM, 25 µM, 50 µM, 100 µM, 200 µM, and 400 µM of purchased sTyr (Bachem) dissolved in methanol-water (2:3) were also prepared and injected with same volume by the same method. Using LC-MS data, a linear standard curve was generated based on peak areas corresponding to sTyr ions and the concentration of sTyr in standard samples. The standard curve was then used to calculate the concentration of sTyr from different cell lysates. The intracellular concentration of sTyr in cells was calculated with the following equation.

$$[sTyr\ intracellular] = \frac{sTyr\ concentration\ in\ lysate\ \times volumn\ of\ lysate}{total\ cell\ numbers\ \times\ cell\ volumn}$$

2 pL was used as an average volume of mammalian cells.

**Mass Spectra Methods For Proteins**

A single quadrupole mass spectrometer (Agilent: G7129A) coupled with 1260 infinity II Quaternary Pump (Agilent: G7111B) was used for all the protein samples with PLRP-S (1000A, 5 µm) column. Water with 0.1% formic acid and ACN with 0.1% formic acid were the organic and aqueous mobile phase, respectively. Flow gradient was initially set at 5% ACN, 15% ACN at 0.1 min, 55% ACN at 4.5 min and then back to 10% ACN at 5 min. Spectra were deconvoluted using the Maximum Entropy deconvolution algorithm in the software BioConfirm.
